# Alternative MyD88 -Cyclin D1 signaling in breast cancer cells regulates TLR3 mediated cell proliferation

**DOI:** 10.1101/2020.04.12.037986

**Authors:** Aradhana Singh, Ranjitsinh Devkar, Anupam Basu

## Abstract

TLR3 mediated apoptotic changes in cancer cells are well documented and hence several synthetic ligands of TLR3 are being used for adjuvant therapy. But there are reports showing contradictory effect of TLR3 signaling which includes our previous report that had shown cell proliferation following surface localization of TLR 3. However, the underlying mechanism of cell surface localization of TLR3 and subsequent cell proliferation lacks clarity. This study addresses TLR3 ligand mediated signaling cascade that regulates a proliferative effect in breast cancer cells (MDA MB 231 and T47D) challenged with TLR3 ligand in the presence of MyD88 inhibitor. Evidences were obtained using immunoblotting, co-immunoprecipitation, confocal microscopy, Immunocytochemistry, ELISA, and flowcytometry. Results had revealed that TLR3 ligand treatment significantly enhanced breast cancer cell proliferation marked by an upregulated expression of cyclinD1 but the same were suppressed by addition of MyD88 inhibitor. Also, expression of IRAK1-TRAF6-TAK1 were altered in the given TLR3-signaling pathway. Inhibition of MyD88 disrupted the downstream adaptor complex and mediated signaling through TLR3-MyD88-NF-κB (p65)-IL6-Cyclin D1 pathway. TLR3 mediated alternative signaling of the TLR3-MyD88-IRAK1-TRAF6-TAK1-TAB1-NF-κB axis leads to upregulation of IL6 and cyclinD1. This response is hypothesized to be via the MyD88 gateway that culminates in proliferation of breast cancer cells. Overall, this study provides first comprehensive evidence on involvement of canonical signaling of TLR3 using MyD88 - Cyclin D1 mediated breast cancer cell proliferation. The findings elucidated herein will provide valuable insights into understand the TLR3 mediated adjuvant therapy in cancer.

## 1. Introduction

Toll-like receptor 3 (TLR3) recognizes double-stranded RNA (dsRNA) of viral origin, small interfering RNAs, and self-RNAs derived from damaged cells (Bugge et al., 2017; Kawasaki and Kawai 2014). TLR3 induces a potent antigen-specific CD8+ T-cell responses that directly induces effector CD8+ T-cell and natural killer (NK) cells for IFN-γ release (Conforti et al., 2010). TLR3 was reported to be expressed not only by immune cells but also in the various cancer cells, such as breast cancers (Salaun et al., 2006), prostate cancer (Gambara et al., 2014), epithelial adenocarcinoma (Helminen et al., 2016), and others. Classically TLR3 signaling is mediated through the endosomal compartment of the cells. Intracellular TLR3 signaling can directly induce apoptosis (Conforti et al., 2010; Salaun et al., 2006). TLR3 structure is comprising of the leucine-rich repeat domain, a transmembrane region, a linker region, and a Toll/IL-1 receptor (TIR) domain (Choe 2005; Takeda and Akira 2004). Ligand binding is mediated by the leucine-rich domain, whereas intracellular signaling is propagated by the TIR domain (Conforti et al., 2010; Salaun et al., 2006).

Canonical TLR signaling has been reported to be regulated by an array of molecules through various mechanisms to adjust the consequences of associated autoimmune and inflammatory diseases. In the canonical pathway, for most of the TLRs, upon ligand activation, MyD88 recruited as a dimer in the cytoplasmic TIR domain in a homophilic interaction (Chen et al., 2018; Loiarro et al., 2010; Noursadeghi et al., 2008; Jia et al., 2014; Hsiao et al., 2014; Han et al., 2002).

TLR3 agonists have been used in immunotherapy for various clinical and preclinical studies. The majority of clinical studies establish TLR3 as a tumor suppressor using synthetic ligand poly(I:C) or poly-ICLC for adjuvant therapy or targeted therapy (Jia and Wang 2015; Braunstein et al., 2018; Ho et al., 2015, Schau et al., 2019). Ligand binding has been reported to induce endosomal TLR3 mediated recruitment of TIR domain-containing adapter-inducing interferon β (TRIF) (O’Neill and Bowie 2007) to trigger type-I IFN and to induce cellular apoptosis (Conforti et al., 2010; Salaun et al., 2006; Gambara et al., 2014; Oshiumi et al., 2003; Yamamoto 2003). On the contrary, TLR3 has been reported to be highly expressed in breast tumors and is associated with poor prognosis of the disease (González-Reyes et al., 2010; Jia et al., 2014; Odalys et al., 2019).

Induction of cell proliferation via surface localization of TLR3 has been shown by our research group in breast cancer cells (Bondhopadhyay et al., 2014) and by other groups in diverse types of cancers (Glavan and Pavelic 2014; Bugge et al., 2017). This mechanism is supposedly an alternative to the endosomal mediated action but the exact mechanism of alternating TLR3 signaling lacks clarity. In this study, we have addressed the alternative signaling, independent of TRIF activation to decipher the mechanistic cascade of alternative cellular proliferative mode of TLR3 signaling.

## 2. Materials and Methods

### 2.1. Cell lines and cell culture conditions

Human breast cancer cells MDA-MB-231, and T47D were obtained from National Center for Cell Science, Pune, India. MDA-MB-231 cells were cultivated in L-15 medium (Himedia, India) and T47D cells were grown in RPMI 1640. All the media were supplemented with 10% FBS (GIBCO) and 1% L-Glutamine-Penicillin-Streptomycin (200mM L-Glutamine, 10,000 units/mL Penicillin and 10mg/mL Streptomycin) (Himedia, India). T47D cells were maintained at 37 □C in a humidified incubator with 5% CO_2_ while MDA-MB-231cells were maintained at 37□C in a humidified incubator without CO_2_.

### 2.2. TLR3 Ligand

Poly(I:C) HMW (Invivogen, Catalog # tlrl-pic) was used as synthetic ligand of TLR3. Accordingly, a dose of 10 μg/ml of Poly(I:C) was used in serum free culture media to bind with TLR3 present in cell.

### 2.3. MyD88 inhibitor

MyD88 inhibitor ST2825 (MCE - HY-50937) was used to block the dimerization of MyD88. Cells were treated with ST2825 (1 μM), for 4 hours prior to addition of poly(I:C).

### 2.4. Cell proliferation assay

Breast cancer cells were seeded in 35 mm culture dish at a density of 40×10^4^ cells in appropriate culture media supplemented with 10% FBS and allowed to grow for 24 hours. After the cells have reached nearly 40-50% confluence, cells were treated with MyD88 inhibitor, 4 hours prior to addition of TLR3 ligand in serum-free media. At the end of incubation period, cells were trypsinized and stained with trypan blue and counted the number of viable cells under microscope. Two technical replicates per sample were run in each independent experiment.

### 2.5. BrdU Incorporation assay

To confirm active DNA synthesis as confirmatory index of cellular proliferation, BrdU incorporation assay was carried out through flow cytometry. Briefly, cells were plated in 35 mm culture dish at 40×10^4^ cells per dish and allowed to adhere overnight in complete media at 37^0^C and treated with MyD88 inhibitor, 4 hours prior to addition of TLR3 ligand. At the end of culture, 10uM BrdU (BD Pharmingen BrdU Flow Kit, San Diego, CA, USA) was added and the target cells were incubated for another 30 minutes, the medium was discarded and the cells were fixed at room temperature for 30 minutes. Cells were permeabilized and FITC conjugate anti-BrdU antibody (BD Pharmingen), was allowed to bind with the incorporated BrdU. After washing, cells were incubated with 7AAD and acquired through BD FACSVerse flow cytometer (BD Biosciences, San Diego, CA, USA).

### 2.6. Immunocytochemistry

To check expression of TLR3 in cell surface as well as the level of IL-6 in the cytoplasm, immunocytochemistry was performed. Briefly, 40×10^4^ cells were seeded on cover slip in 35mm culture dish in complete media. Cells were allowed to adhere for overnight and treated. Four hours before the addition of TLR3 ligand, MyD88 inhibitor was added and incubated for 24 hours. For TLR3 surface expression, cells were fixed and allowed to bind with TLR 3 antibody (Invitrogen-PA5-29619) and Alexa 594 conjugated secondary anti-rabbit goat antibody (Invitrogen-A11012). For IL-6 expression Cells were fixed, permeabilized and incubated with primary IL-6 antibody (Invitrogen-AMC0862) and Alexa 488 conjugated anti-mouse goat antibody (Invitrogen-A11001) and mounted with Vecta Shield – DAPI to counter stain nuclei and observed under fluorescence microscope (Leica DMI 6000B).

### 2.7. Confocal microscopy study

Confocal microscopy was carried to study the change in nuclear localization of NF-κB in breast cancer cells after TLR3 ligand activation. Cells were seeded with a density of 40×10^4^ on coverslip with 35 mm culture plate. Cells were treated with TLR3 ligand with or without MyD88 inhibitor for 30 minutes, 60 minutes and 90 minutes. Cells were fixed with 3% PFA (paraformaldehyde solution) for 15 minutes at room temperature, washed with PBS and transferred to 100% methanol for 5 minutes, washed with PBS and permeabilized with PBS containing 0.25% Triton X-100 for 5 min. After fixation and permeabilization, blocking was done using PBS containing 1% BSA for 1 hours. After blocking, cells were allowed to bind with NF-κB p65antibody (Invitrogen-PA1-186) for overnight at 4°C followed by anti-rabbit antibody conjugated with Alexa-594 for 1 hours at room temperature in dark. Coverslips were mounted with Vecta Shield-DAPI to counterstain nuclei and analyzed by Zeiss LSM 710 inverted confocal microscope (Zeiss, Germany) with an 63X plan apochromat objective. Image analysis was performed using ImageJ v3.91 software (http://rsb.info.nih.gov/ij).

### 2.8. ELISA

IL-6 was quantified from the cell supernatant of the challenged cells by ELISA. Briefly, cells were seeded at a density of 100×10^4^ cells in 35 mm culture plate and grown to confluence. Confluent monolayer was washed twice and kept in media supplemented with 1% ITS. Then incubation with TLR3 ligand and MyD88 inhibitor was performed for 36 hours. Two replicates per sample were run in each independent experiment. At the end of incubation, condition media was collected and estimated quantity of the secretary IL-6 using commercially available ELISA kit (R&D Systems, DY 206-05) with human IL6 antibody (R&D Systems, DY 008).

### 2.9. Western blotting

Cells were cultured at a density of 100×10^4^ cells in 60 mm culture plate (Tarsons-960020). Four hours before the addition of TLR3 ligand, MyD88 inhibitor was added and incubated for 24 hours. Cells were lysed using RIPA buffer and gel electrophoresis was performed using acrylamide gel. Proteins were transferred to PVDF Membranes and blotted with antibodies against IRAK1 (Invitrogen-38-5600), phospho IRAK1-Thr209, (Invitrogen-PA5-38633), TAK1 (Invitrogen-700 113), phospho TAK1-Thr184/187 (Invitrogen-MA5-15073), TAB1 (Invitrogen-PA5-28683), TRAF-6 (Invitrogen-PA5-29622), and Cyclin D1 (Invitrogen-AHF0082). The antibody against β-actin (Invitrogen-MA191399) was used as a loading control.

### 2.10. Co-immunoprecipitation

Cells were cultured at a density of 100×10^4^ cells in 60 mm culture plate (Tarsons-960020). Four hours before the addition of TLR3 ligand, MyD88 inhibitor ST2825 was added and incubated for 24 hours. Cells were lysed with non-denaturing lysis buffer (20mM Tris-HCl, 137mM NaCl, 1% Triton X-100, 2mM EDTA with protease inhibitor cocktail. The lysate was incubated on ice for 30 minutes, and centrifuged at 10,000 rpm for 20 minutes at 4^0^C. Supernatant were incubated with 1μg of indicated antibody and dynabeads (Invitrogen-10003D) for overnight at 4^0^C. The dynabeads were pellet down and washed with lysis buffer after overnight incubation. The precipitates were resolved in SDS-PAGE and subjected to western blotting with the indicated antibodies.

### 2.11. Statistical analysis

Statistical analysis was performed with GraphPad Prism version 7. The difference between two groups were determined by two-tailed Student’s T - test. Two or more groups were compared with one-way ANOVA. A p-value <0.05 was considered for statistically significant.

## 3. Results

### 3.1. TLR3 ligand induces cell proliferation which is restricted by the MyD88 inhibitor

To verify the alternative TLR3 signaling, breast cancer cells were pretreated with MyD88 inhibitor (ST2825) for 4 hours followed by stimulation of TLR3 by addition of the TLR3 ligand. This was followed by incubation of the cells for 24 hours. We have observed a significant increase in cell proliferation of MDA-MB-231 and T47D cells. The MyD88 inhibitor impaired the proliferative effect of TLR3 ligand poly(I:C) in both MDA-MB-231 and T47D cells (Fig. 1A-B).

**Fig 1.**
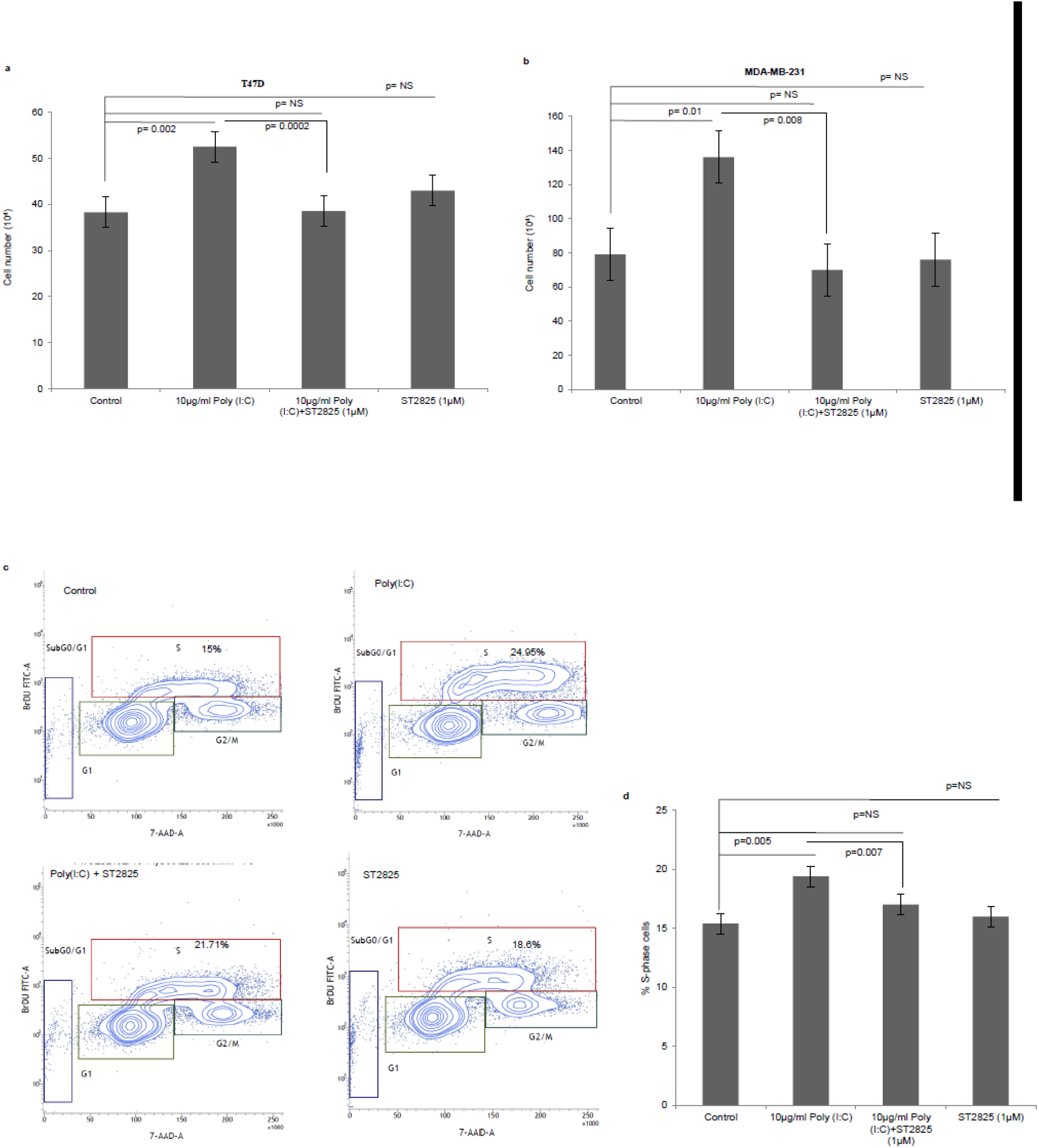
TLR3 ligand induces cell proliferation and stunted by MyD88 inhibitor. Breast cancer cells were pre-treated with MyD88 inhibitor (1μM) 4 hours prior to addition TLR3 ligand (10μg/ml) for 24 hours before cells were counted. Control indicates cells were not treated with TLR3 ligand or MyD88 inhibitor. Growth kinetic assay. **(A)** T47D and **(B)** MDA-MB-231 showing proliferative effect of TLR3 ligand. Inhibition of MyD88 dimerization restrict the proliferative effect. **(C)** Contour plots for BrdU – Cell proliferation assay using T47D cells. Cells were pretreated with MyD88 inhibitor (1μM) 4 hours prior to addition TLR3 ligand (10μg/ml) for 24 hours before labelled with BrdU and detected by flow cytometry. Contour plots of DNA -7AAD-A versus log BrdU-FITC showing G_0/_G_1_, S and G2/M gates of Control cells, Cell treated with TLR3 ligand, cells treated with TLR3 ligand and MyD88 inhibitor and cells treated with only MyD88 inhibitor. **(D)** Bar graph showing percentage of S-phase gated cells among the different experimental cell groups following BrdU incorporation. The results are presented as mean ± S.D and p< 0.05 is treated as significant).

Further to confirm cellular proliferation, BrdU incorporation assay was undertaken that revealed higher percentage of BrdU incorporated in S-phase cells in TLR3 ligand treated cells as compared to untreated cells. The proliferative effect of TLR3 ligand treatment was nullified by treatment with the MyD88 inhibitor (Fig. 1C-D). There was no cytotoxic or cell proliferating effect has been observed in the cells that has been treated with only MyD88 inhibitor.

### 3.2. TLR3 ligands stimulates the expression of surface TLR3

To confirm our previous findings of expression of TLR3 on the surface breast cancer cells (Bondhopadhyay et al., 2014), in this study we had verified the membrane expression of TLR3 in MDA-MB-231 and T47D breast cancer cells in the absence or presence of exogenous TLR3 ligand and MyD88 inhibitor by immunocytochemistry. TLR3 expression was markedly increased in the presence of exogenous TLR3 ligand in comparison to the unstimulated cells, while the addition of MyD88 did not effect on TLR3 expression (Fig 2).

**Fig 2.**
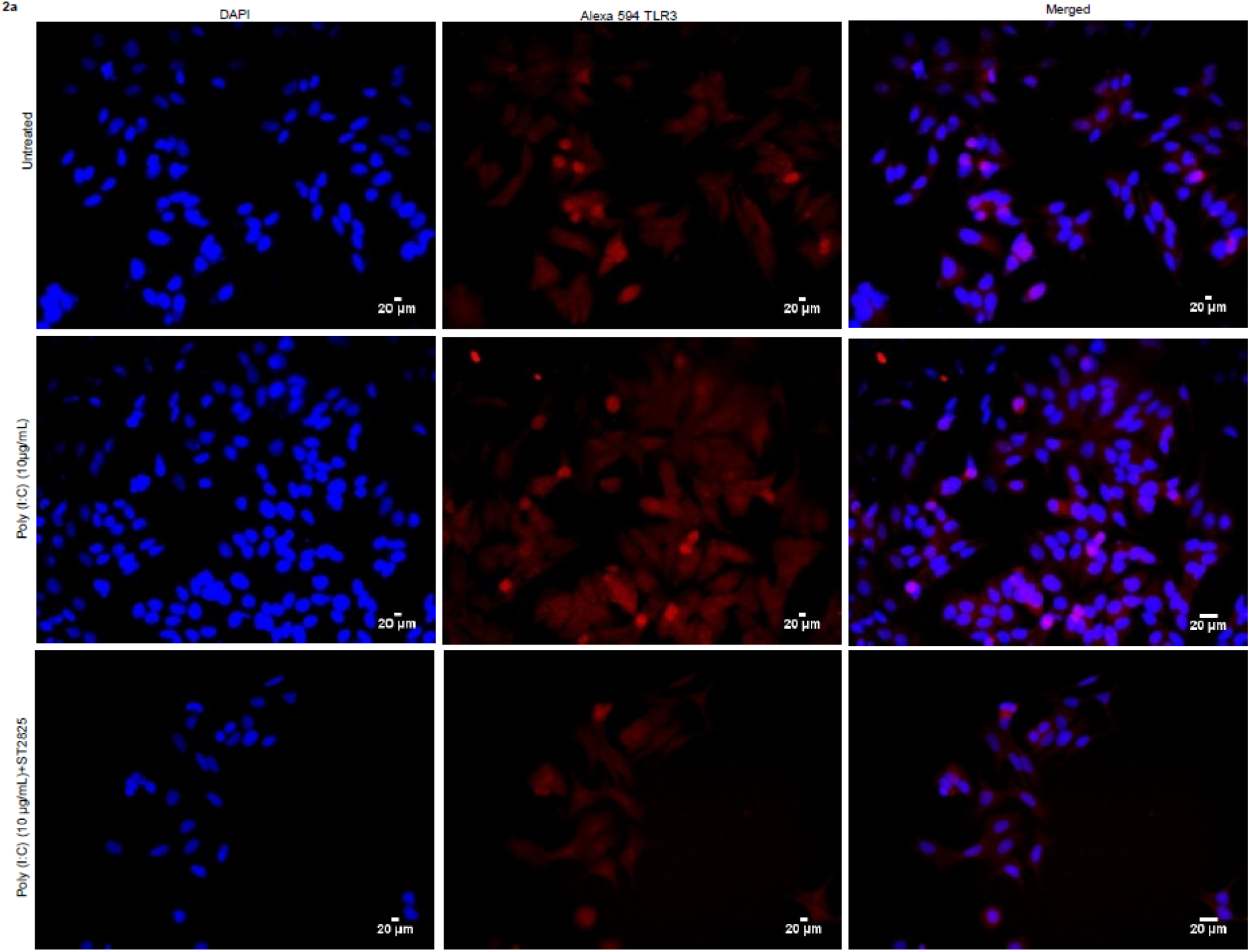
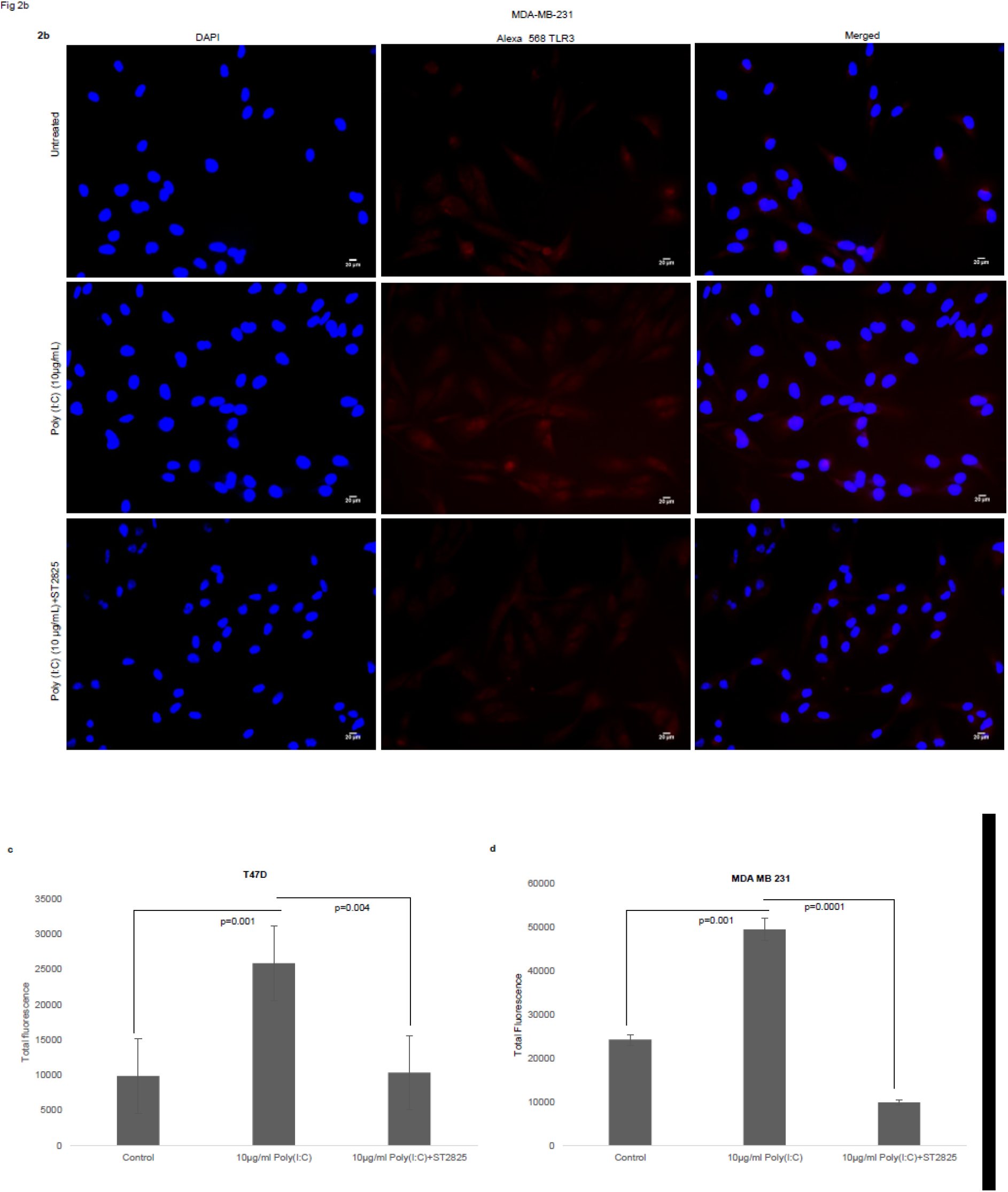
Effect of MyD88 inhibitor on surface localization of TLR3. Fluorescent microscopy image of cells, treated with TLR3 ligand (10μg/ml) with or without MyD88 inhibitor (1μM) following immunocytochemical staining with antibody against TLR3 and Alexa 594 tagged secondary antibody and counterstained with DAPI. **(A)** T47D cell **(B)** MDA MB 231 Cells. **(C) and (D)** Bar graph showing the localization of TLR3 in cells surface after observe through the microscope and analyse through the ImageJ package for all the experiment groups. The results are presented as mean ± S.D (p< 0.05 is treated as significant).

### 3.3. MyD88 inhibitor reduces the production of proinflammatory cytokine IL-6 in TLR3 ligand treated breast cancer cells

To assess, whether TLR3 ligand able to induce IL-6 production and be reversed, cells were treated with MyD88 inhibitor 4 hours. Accordingly, MyD88 inhibitor pretreated cells were challenged with TLR3 ligand and level of IL6 was determined by immunocytochemistry in cytoplasm and by ELISA in condition media. We have observed that TLR3 ligand treatment significantly induces the immunofluorescence and secretion of IL-6 compared to the control group. Pretreatment of MyD88 inhibitor reduced the production of IL-6 in spite of stimulation with TLR3 ligand (Fig. 3).

**Fig 3.**
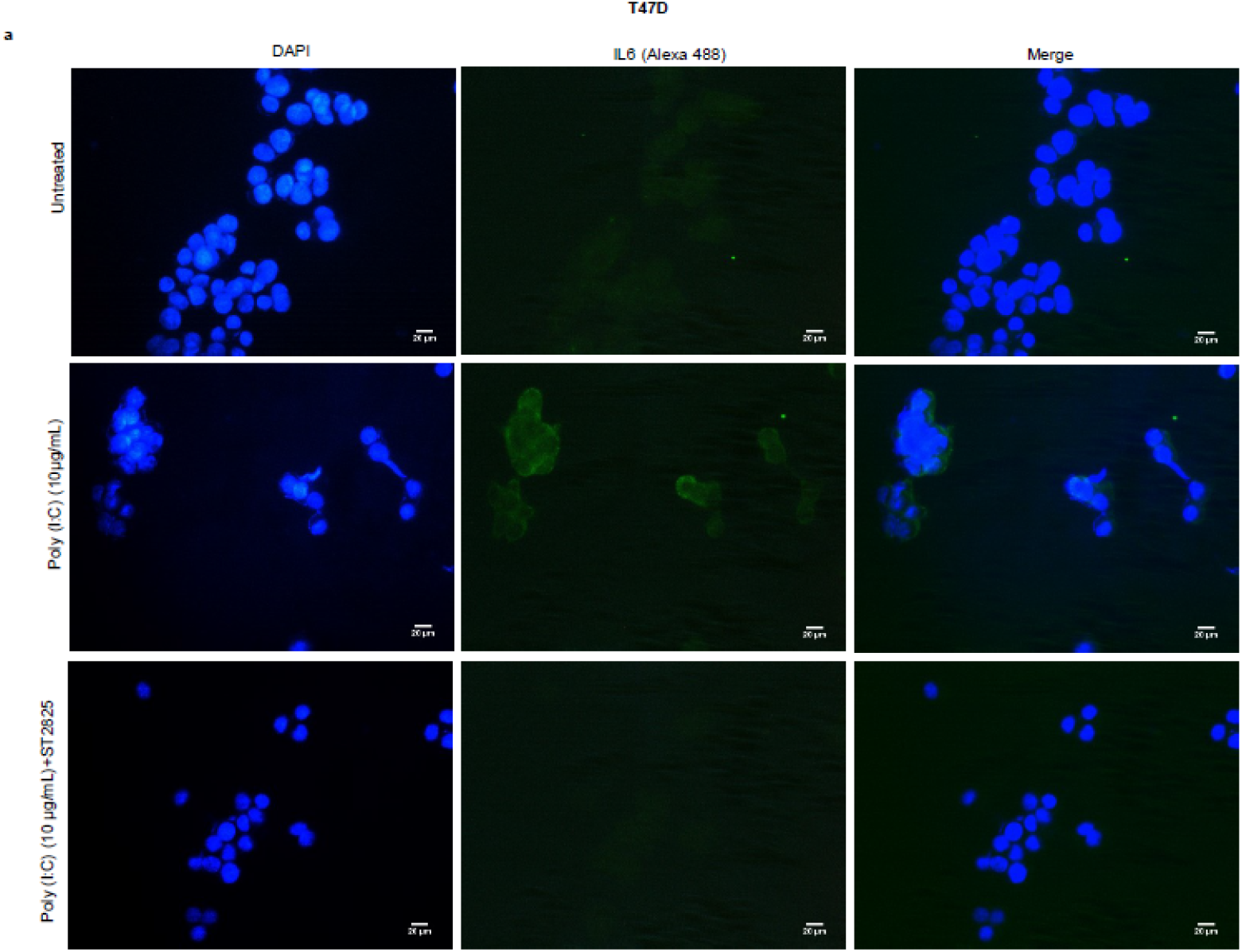
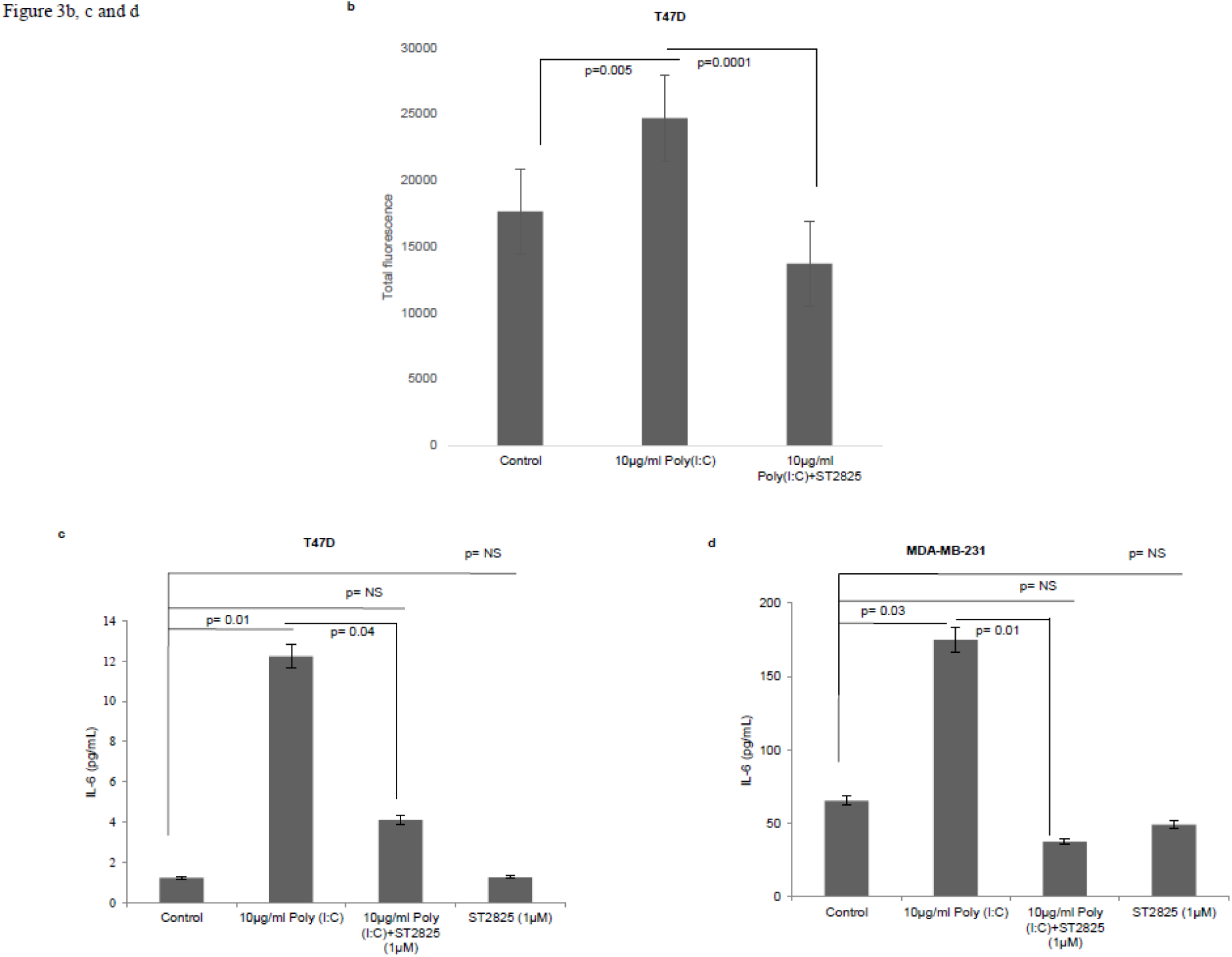
Expression of IL-6 following MyD88 inhibitor and TLR3 ligand treatment. **(A)** Fluorescent microscopy image of T47D cells, treated with TLR3 ligand (10μg/ml) with or without MyD88 inhibitor (1μM) following immunocytochemical staining with antibody against IL-6 and Alexa 488 tagged secondary antibody and counterstained with DAPI. Untreated indicates the cells are not treated with TLR3 ligand. (magnification, 40X). **(B)** Bar graph showing the expression of IL6 following observe through the microscope and analyse through the ImageJ software for all the experiment groups. **(C), (D)** Expression of IL6 in the cell culture supernatant as measured through the ELISA. The results are presented as mean ± S.D (p< 0.05 is treated as significant).

### 3.4 MyD88 inhibitor attenuates TLR3 ligand-induced NF-κB nuclear localization

In the previous section we have showed that there was reduction in the IL-6 expression after addition of MyD88 inhibitor despite the presence of TLR3 ligand. Previously it was reported that early phase activation (0.5-2h) of NF-κB leads to the production of pro-inflammatory cytokines (Han et al., 2002). In the present work, we had assessed the early phase nuclear localization of p65 subunit of NF-κB. Accordingly, it has been observed that MyD88 inhibitor nullifies TLR3 ligand induced nuclear localization of p65 (Fig 4). TLR3 ligand elicits highest translocation of p65 into the nucleus at the 60-minute time point in both the cell lines compared to control untreated cells (Supplementary Figure 1).

**Fig 4.**
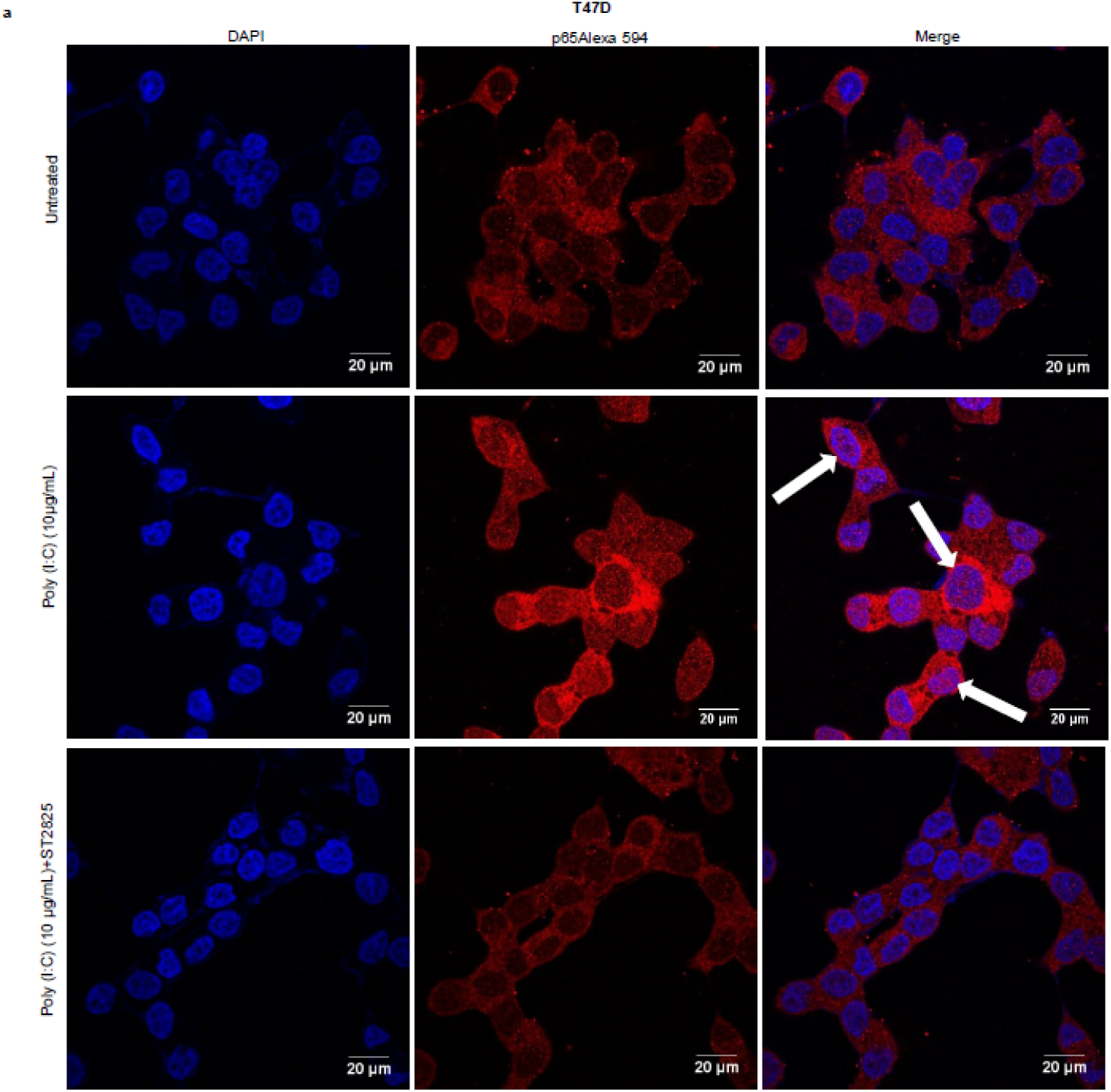
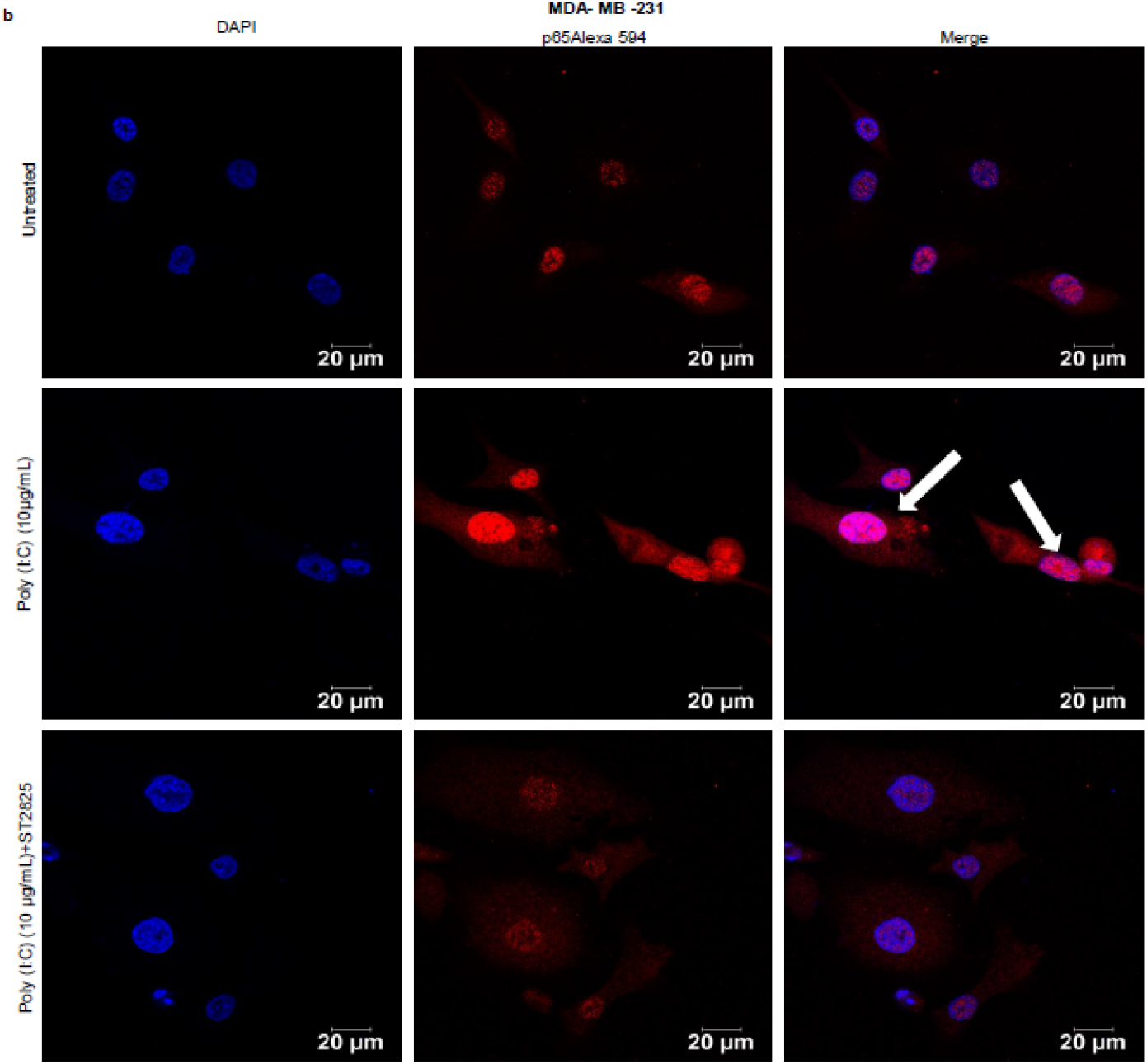
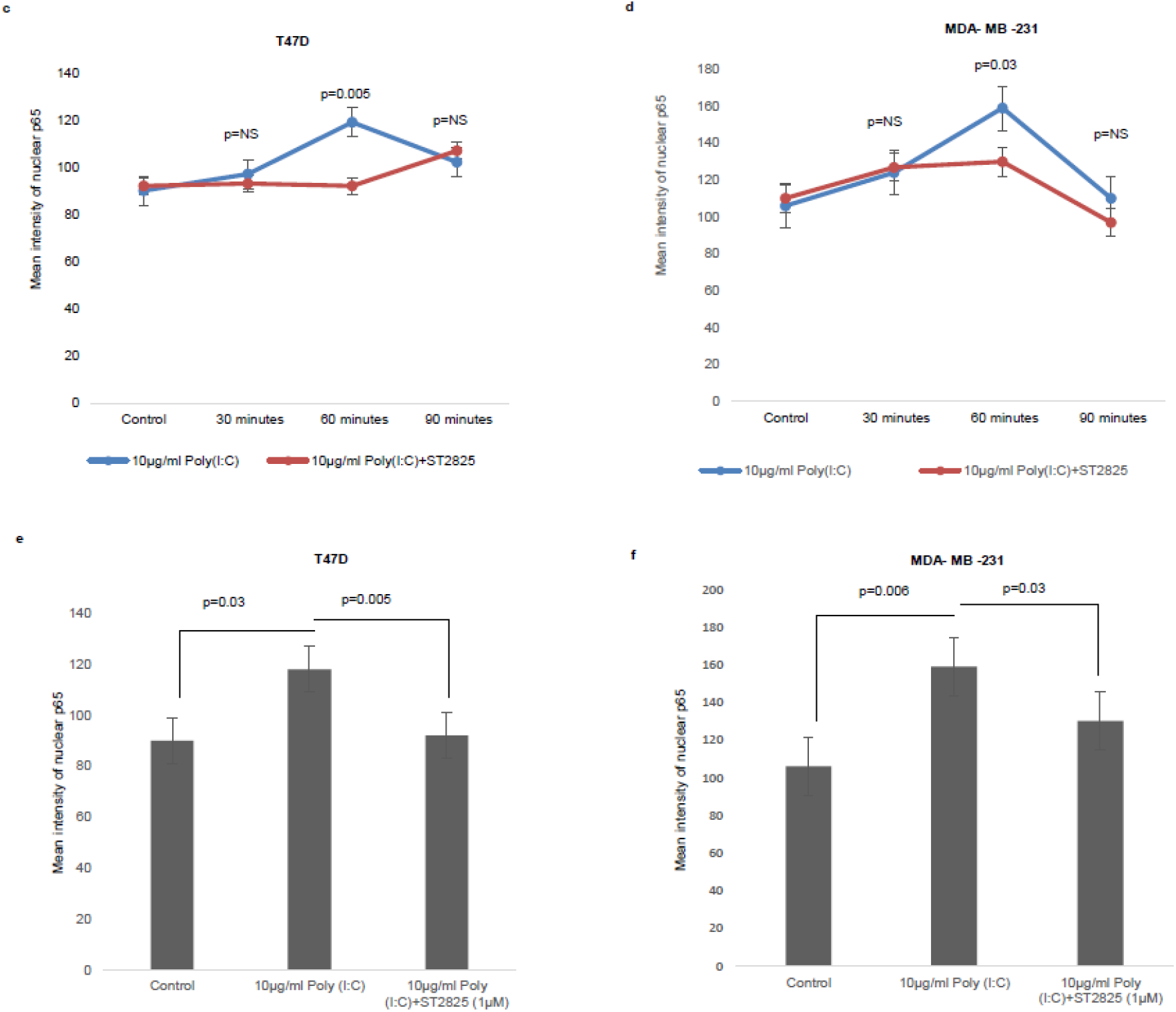
Confocal microscopy for nuclear translocation of p65. **(A)** T47D cells, **(B)** MDA-MB-231 cells were pre-treated with MyD88 inhibitor for 4 hours prior to addition of TLR3 ligand (10μg/mL) for 60 minutes. Cells were stained with antibody against p65 subunit of NF-κB and Alexa 594 tagged secondary antibody and counterstained with DAPI and image acquired through confocal microscope (magnification, 63X); **(C) and (D)** Bar graph is presented as mean ± S.D for the quantitative measurements of nuclear localization of NF-κB at 30 minutes, 60 minutes, 90 minutes of stimulation, analysed through Image J package. (p< 0.05 is treated as significant). e,f Bar graph at 60 minutes of stimulation showing the highest nuclear localization of NF-κB.

#### TLR3 ligand induced the expression of Cyclin D1 and halted by the MyD88 inhibitor

To address that TLR3 mediated cell proliferation, whether regulated through the cyclin D1 gene expression, we investigated the expression of cytosolic cyclin D1 through western blotting using cell lysate. TLR3 ligand stimulation elevates the expression of cyclin D1. However, the addition of MyD88 inhibitor recorded a decrement in the level of cyclin D1 suggesting a break in the signaling cascade of TLR3 ligand (Fig. 5 A-B).

**Fig 5.**
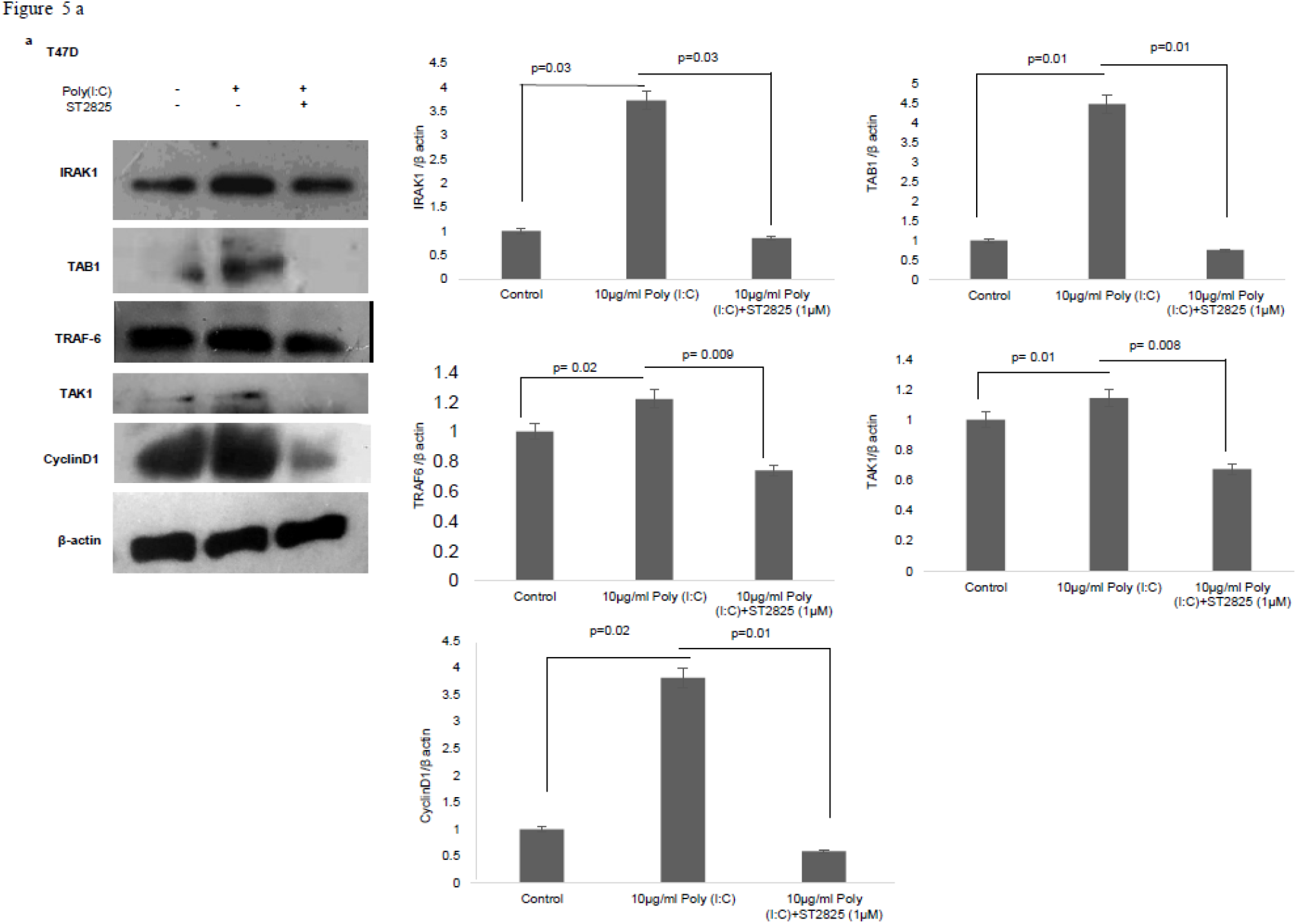
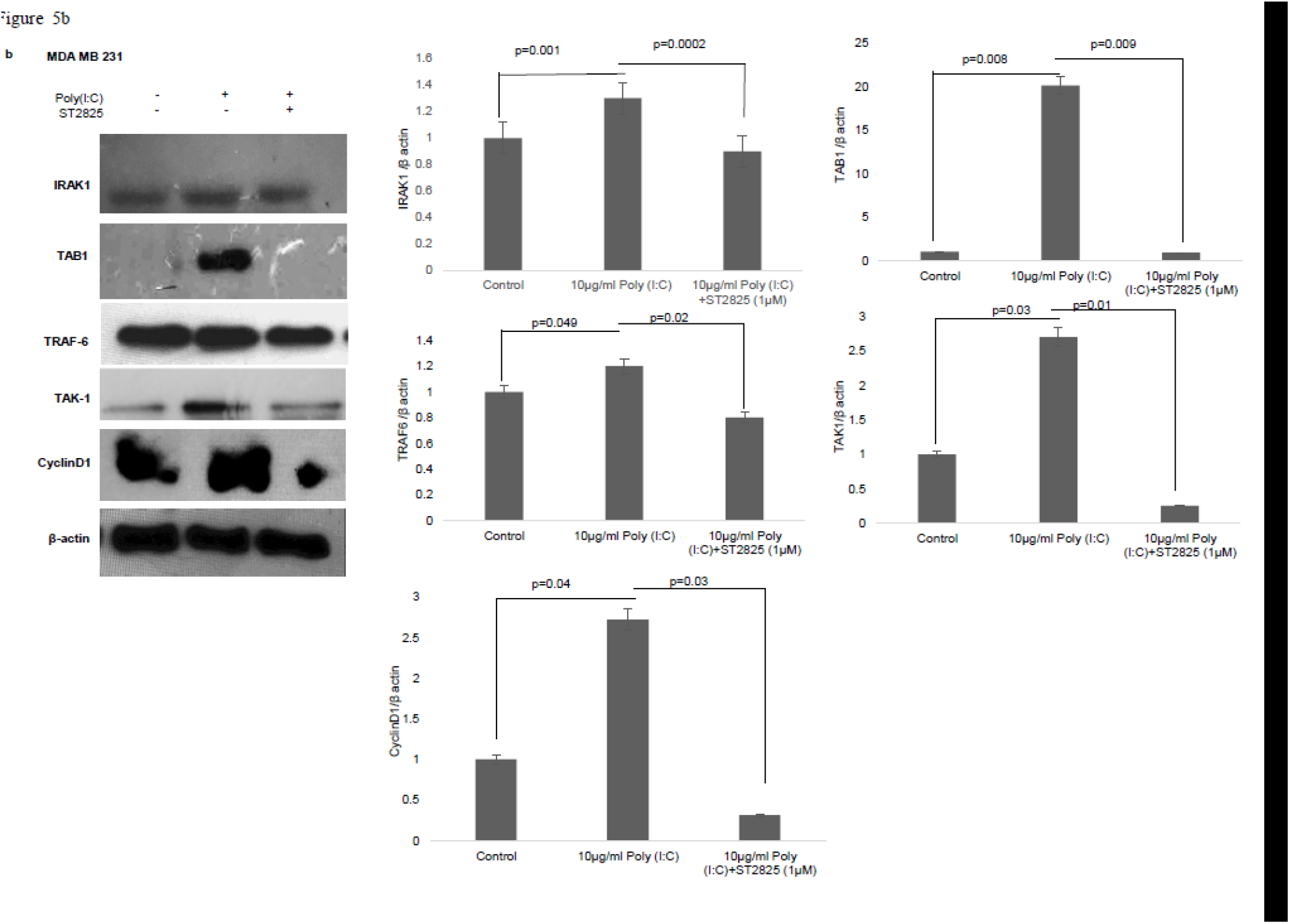
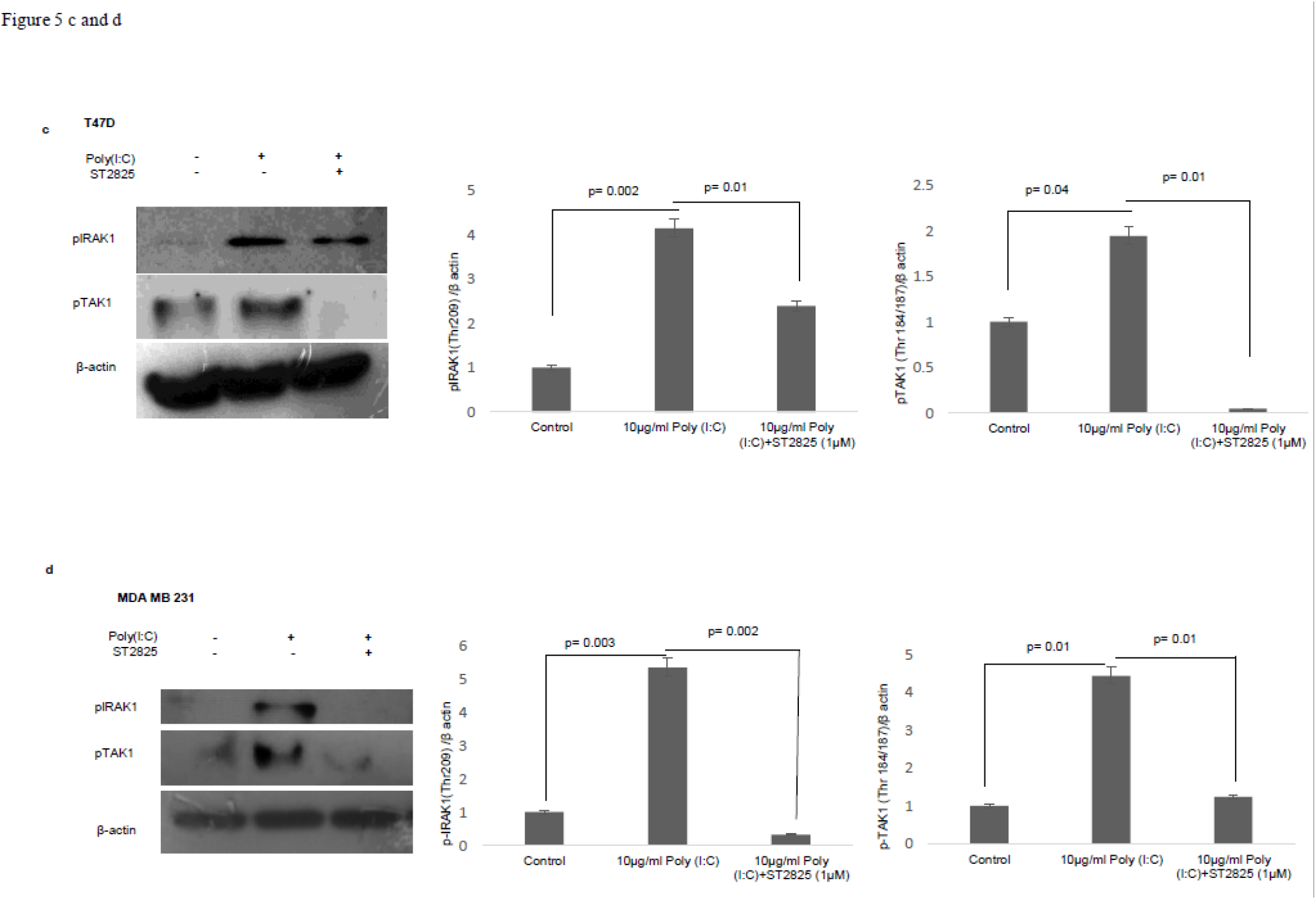
Western-blotting for the expression of signalling protein. **(A)** and **(B)** Cell lysate were collected and subjected to western blot assay to estimate the level of the expression of IRAK1, TAK1, TAB1, TRAF-6 and cyclin D1. **(C) and (D)** Expression of pIRAK1 and pTAK1. β-actin was used as loading control. The respective bar graphs are presented as densitometry analysis as mean ± S.D of experiments (p < 0.05 is treated as significant).

#### MyD88 inhibitor reduced the exogenous TLR3 ligand -induced expression of adaptor proteins -IRAK1, TAK1, TAB1 and TRAF6

To understand the involvement adopter complex to transmit the effect of TLR3 ligand for the expression of Cyclin D1, we have checked the expression IRAK1, TRAF6, TAK1, TAB1 in the presence or absence of the MyD88 inhibitor. To confirm our hypothesis, protein level of all the above adaptor proteins has been estimated by western blotting. The significant increase in expression of IRAK1, TRAF6, TAK1, and TAB1 following induction of TLR3 ligand has been observed compared to untreated cells. However, addition of MyD88 inhibitor ST2825 reduced the level of IRAK1, TRAF6, TAK1, and TAB1(Fig 5A-B).

#### MyD88 inhibitor reduces TLR3 ligand mediated phosphorylation of adaptor protein IRAK1 and TAK1

IRAK1, a serine-threonine kinase, was reported to be phosphorylated upon lipopolysaccharide (LPS) mediated signaling stimulation (Dong et al., 2006). We, have assessed IRAK-1 and TAK1 phosphorylation in MDA-MB-231 and T47D cells in presence and absence of MyD88 inhibitor following the induction by TLR3 ligand. Increased in the level of phosphorylated IRAK1 and TAK1 in the response of exogenous TLR3 ligand addition potentially explain the activation of IRAK1 and TAK1. We found that MyD88 inhibitor suppressed the TLR3 ligand mediated level of phosphorylated IRAK1 and TAK1 (Fig. 5 C-D).

### 3.4. TLR3 ligand induces IRAK1/TRAF6, p-IRAK1 /TAK1 and TRAF6/TAK1/TAB1 interactions which are disrupted by MyD88 inhibitor

As there were change of expression and phosphorylation, further we have addressed the involvement of signaling complex formation of the above adopter proteins. Accordingly, cells were treated with TLR3 ligand in presence or absence of MyD88 inhibitor, thereafter immunoprecipitated with IRAK1 antibody and immunoblotted with TRAF6 antibody were investigated. In TLR3 ligand stimulated cells there was a marked increase in association. In cells pretreated with MyD88 inhibitor before TLR3 ligand addition, the interaction of TRAF6 and IRAK1 was decreased markedly. This suggests that MyD88 inhibition interferes with the formation of the TLR3 ligand-induced IRAK1/TRAF6 complex. (Figure 6A).

**Fig 6.**
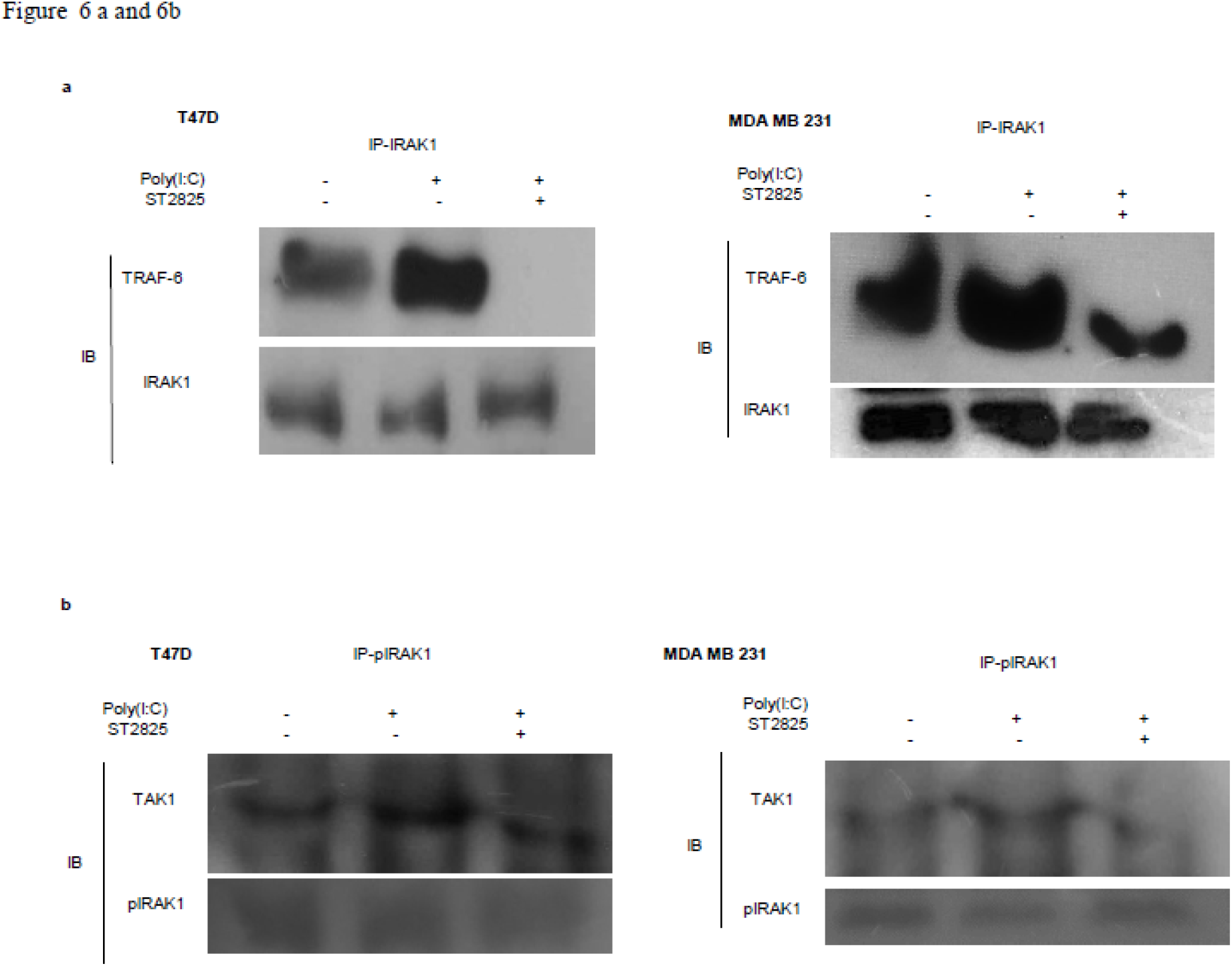
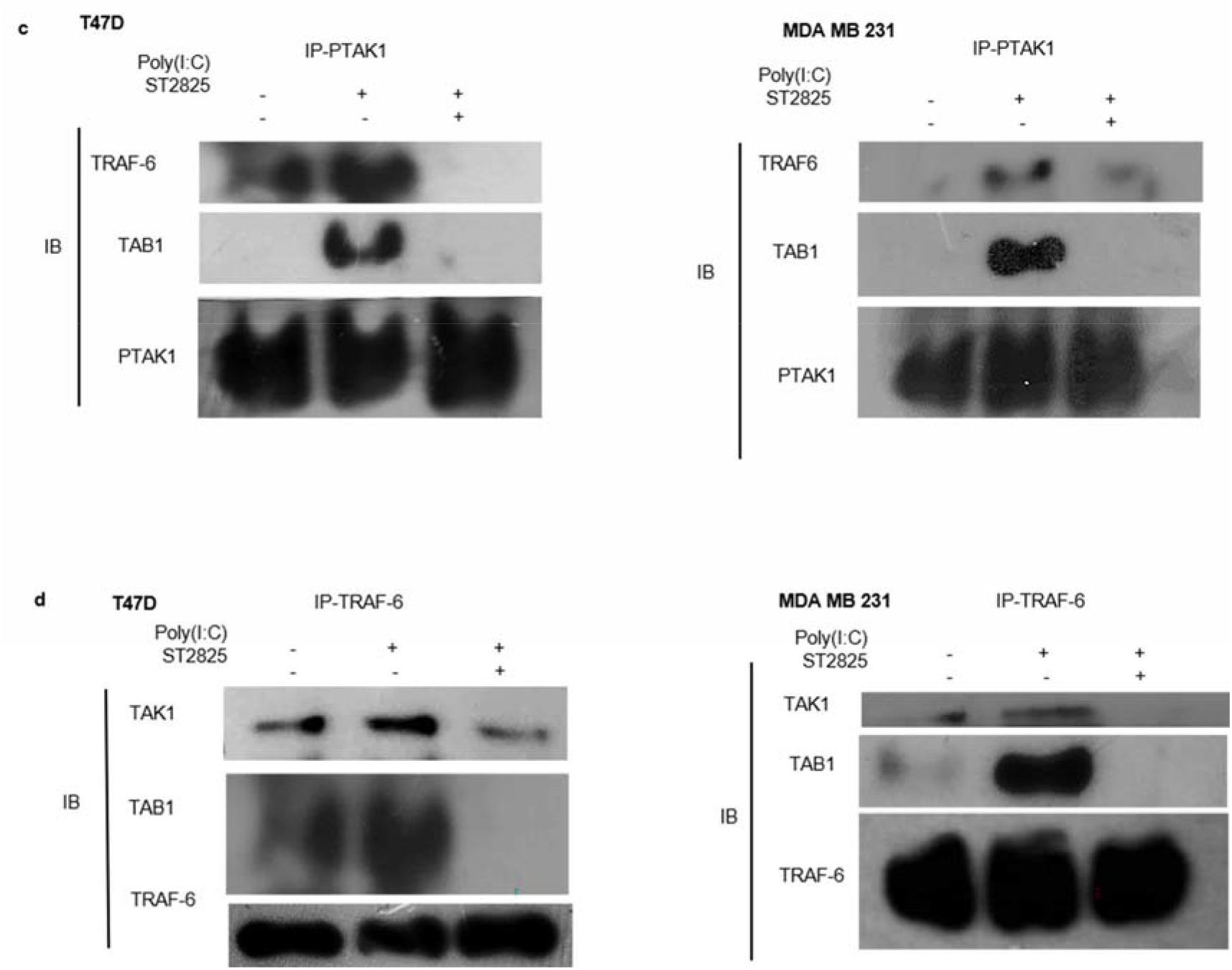
Immunoprecipitation showing the involvement of the signalling complex **(A)** signalling complex of IRAK1/ TRAF-6 was immunoprecipitated with antibodies against IRAK1 followed by western blotting with anti-TRAF-6 and anti-IRAK1 antibody. **(B)** signalling complex of pIRAK1/ TAK1 was immunoprecipitated with antibodies against pIRAK1 followed by western blotting with anti-TAK1 and anti-pIRAK1 antibody. **(C)** signalling complex TAB1-TRAF6-TAK1 was immunoprecipitated with antibodies against pTAK1 followed by western blotting using anti-TRAF6, TAB1 and pTAK1 antibody **(D)** signalling complex TAB1-TRAF6-TAK1 was immunoprecipitated with antibodies against TRAF6 followed by western blotting analysis using anti-TAK1, TAB1 and TRAF6 antibody.

On the other hand, cell lysate was immunoprecipitated with pIRAK1 antibody and immunoblotted with TAK1 antibody. In TLR3 ligand stimulated cells there was a marked increased association. In cells pretreated with MyD88 inhibitor, the interaction of TAK1 with Phospho-IRAK1 was decreased markedly. This suggests that MyD88 inhibitor ST2825 interferes with the association of TLR3 ligand induced complex of pIRAK1/TAK1 (Figure 6B). It is also worthy to mention that we did not find any immunoprecipitation of TAK1 when precipitated thorough non phosphorylated IRAK1 antibody. As we had mentioned in Fig. 5A and 5B, inhibition of MyD88 dimerization, block the proinflammatory signaling and lower the level of TRAF6, TAK1, and TAB1. Herein, we hypothesize that MyD88 inhibitor interferes with the formation of TLR3 ligand induced MyD88 mediated TRAF6/TAK1/TAB1 signaling complexes. To evaluate this hypothesis, cells were pretreated with MyD88 inhibitor ST2825 for 4 hours prior to stimulation by TLR3 ligand. Cell lysates were collected and immunoprecipitated with TRAF6 and phosphoTAK1 antibodies, followed by immunoblotting using TAB1, TRAF6, and TAK1 antibodies. In TLR3 ligand stimulated cells, there was a distinct increase in the association; whereas, in the cells pretreated with MyD88 inhibitor, the interaction of TAK1 with either TRAF6 or TAB1 was decreased markedly. This suggests that MyD88 inhibitor (ST2825) interferes with the formation of signaling complex as mentioned above (Fig. 6 C and 6D).

## 4. Discussion

In this present study, we had reported the mechanistic pathway of TLR3 ligand-induced breast cancer cell proliferation through MyD88 mediated gateway. ST2825 is well established as a MyD88 inhibitor in several studies (Kawasaki and Kawai 2014; Deng et al., 2016; Shiratori et al., 2017; Loiarro et al., 2007) and hence, was used to address the ligand-mediated alternative cell proliferative action of TLR3. Care was taken that ST2825 is used at a concentration that is neither cytotoxic nor cell proliferative to the breast cancer cell lines (MDA MB 231 and T47D). TLR3 was reported to be expressed only by immune cells and unstimulated TLR3 mainly resides in ER. Stimulation of TLR3 with ligand poly (I:C) it gets translocated from ER into the endosomal compartment (Johnsen et al., 2006). Though, regulation of this translocation is reported to be controlled by UNC93B1 protein, its inhibition has only a partial effect on TLR3 mediated signaling (Bugge et al., 2017). Further, cell surface expression of TLR3 has been reported in a variety of cells such as pulmonary cells, hepatocytes, breast cancer, prostate cancer and epithelial adenocarcinoma (Salaun et al., 2006; Gambara et al., 2014; Helminen et al., 2016) indicate that it signaling occurs through the plasma membrane. Dynasore, a dynamin inhibitor that inhibits endocytosis of the receptors, only partially affects the poly(I:C) mediated TLR3 signaling. This suggests that TLR3 signaling may occur independent to ligand-internalization (Bugge et al., 2017). Our result shows the increase in surface expression of TLR3 in breast MDA-MB-231 and T47D cells upon TLR3 ligand activation. Addition of MyD88 inhibitor does not have any effect on the level of surface TLR3 expression suggesting that TLR3 signaling occurs from the cell surface in MyD88 dependent manner.

We have shown in our initial experiments that exogenous stimulation of TLR3 by its ligand promotes the cellular proliferation in breast cancer cells (Bondhopadhyay et al., 2015). But this proliferative effect has been perturbed by the addition of MyD88 inhibitor suggesting that the said effect of TLR 3 is mediated by the MyD88. In the present investigation, TLR3 activation through TLR3 ligand stimulated the expression of downstream signaling factors, including IRAK1, TRAF6, TAB1, and TAK1 and suppressed the MyD88 inhibitor that correlates with other signaling cascade (Kong et al., 2017). It has been reported that activation of other TLRs, in contrast to TLR3, can induce canonical pathway through MyD88 mediated activation of Interleukin 1 Receptor Associated Kinase 1(IRAK1). Activation of this signaling pathway further regulates IRAK1 mediated activation of TNF Receptor Associated Factor 6 (TRAF6) and Transforming growth factor beta-activated kinase 1 (TAK1) that further causes activation of TGF-Beta Activated Kinase 1 (TAB1). Thus our findings are well correlated with earlier reports for different convergent signaling pathways (Kong et al., 2017; Dong et al., 2006; Cui et al., 2012; Xiong et al., 2011, Johnsen and Whitehead 2006; Rhyasen and Starczynowski 2014, O’Neill and Bowie 2007, Conforti et al., 2010; Salaun et al., 2006; Gambara et al., 2014; Oshiumi et al., 2003; Yamamoto 2003, Ma et al., 2018, Klein and Assoian 2008, Alt et al., 2000).

Though, MyD88 does not have any catalytic activity, its activation causes dimerization leading to the activation of downstream kinases (Chen et al., 2018). It has been shown that the progression of signaling pathways occur due to the phosphorylation of two key adaptor proteins IRAK1 and TAK1. IRAK1, a serine-threonine kinase, was reported to be phosphorylated via MyD88 upon lipopolysaccharide (LPS) stimulation (Dong et al., 2006) that also triggers its dissociation from the membrane and translocation into cytosol. IRAK1 activation is also required for phosphorylation of TAK1 (Dong et al., 2006). We had observed that activation of IRAK1 leads to its complex formation with TRAF6 and TAK1. Phosphorylation of IRAK1 helps in dissociation of TRAF6 complex from the membrane, and may facilitate formation of TRAF6, TAK1, and TAB1 complex in the cytosol. Later, the phosphorylated IRAK1 may get ubiquitinated and degraded as suggested by other research groups (Kong et al., 2017; Dong et al., 2006; Cui et al., 2012; Xiong et al., 2011). Thus, dissociation of the complex from the membrane may lead to phosphorylation of TAK1, as has been shown by research groups in other signal pathways (Cui et al., 2012). In our study, the level of phosphorylation of IRAK1 and TAK1, as well as the association of signaling complex IRAK1/TRAF6, pIRAK1/TAK1 and TRAF6/TAK1/TAB1, was found to be elevated upon TLR3 induction. But, the level of phosphorylation and as well as the interaction and formation of signaling complexes was found to be reduced by administration of MyD88 inhibitor. These findings indicate the TLR3 act in the TLR3-MyD88-IRAK1-TRAF6-TAK1 axis to promote cellular proliferation.

In recent studies, TAK1 has been identified as a key regulator of various immune responses and inflammatory reactions that promote tumorigenesis, fibrosis, and multiple inflammatory disorders. Accordingly, we have observed that, induction of TAK1 phosphorylation as a MyD88 activation cascade, induces the NF-κB activation followed by secretion of IL-6. This observation is supported by an earlier study wherein inhibition of TAK1 phosphorylation inhibits IL-6 production through an NF-κB dependent manner (Hsiao et al., 2014). As NF-κB is dimer composed of p65 and p50 subunits (McFarland et al., 2013; Yang et al., 2014; Brasier 2010), activation of this TAK1/TAB complex activates NF-κB signaling pathway, which induce nuclear localization of p65 (Brown et al., 2010). In our study, we had recorded an early phase activation of NF-κB that had triggered IL-6 release by TLR3 ligand induction, but had shown a downregulation following inhibition of MyD88. LPS induction in mice leads to biphasic stimulation of NF-κB. In the early phase activation (0.5-2h) the production of pro-inflammatory cytokines, tumor necrosis factor (TNF), and IL-1β is seen whereas, the late phase activation (8-12 h) is associated with expression of cyclooxygenase 2-derived anti-inflammatory prostaglandins and the anti-inflammatory cytokines and transforming growth factor-β1 (Han et al., 2002). Our observation is well correlated with this report wherein, an early phase activation leading to production of pro-inflammatory cytokines is observed herein.

The induction of breast cancer cell lines (MDA MB 231 and T47D) with TLR3 ligand induces cellular proliferation through MyD88 dependent manner via induction of pro-inflammatory cytokine IL-6 and Cyclin D1. The addition of MyD88 inhibitor disrupts the signaling pathway that leads to a decreased level of IL-6 secretion as well as lower in cyclin D1 activation. Cyclin D1 controls cell cycle progression through the G1 phase and G1-to-S transition (Ma et al., 2018). Induction of IL-6 has been reported to stimulate cyclin D1 promoter (Ma et al., 2018). Cyclin D1 has been reported to be induced during MyD88-TRAF-6 and TAK-1 signaling pathway via NF-κB-cyclin D1-STAT 3 pathway (Klein and Assoian 2008) and cyclin D1 exported from nucleus to cytoplasm during S-phase of the cell cycle (Alt et al., 2000). TLR3 ligand stimulation elevates the expression of cyclin D1. However, in the present study, it has been observed that addition of MyD88 inhibitor breaks the signaling cascade of TLR3 ligand and hence a decrease in the level of cyclin D1 was recorded. Earlier it has been reported that an elevated IL-6 level in dsRNA-treated TLR3 positive mice, but not in TLR3 negative tumors (Salaun et al., 2011).

TLR3 synthetic ligands were used with conventional chemotherapies or radiotherapy in clinical trials for the treatment of cancer patients (Braunstein et al., 2018; Aranda et al., 2014; Smith et al., 2018). This reported tumor suppressive and apoptotic effect of TLR3 is achieved predominantly by induction of type I IFN and activation of effector cells, when TLR3 is located within the endosomal compartment (Bugge et al., 2017; Gambar et al., 2014; Braunstein et al., 2018). TLR3 synthetic ligand poly-ICLC with Sorafenib significantly reduces tumor growth, both in-vitro and in-vivo in hepatocellular carcinoma (Ho et al., 2015). The mechanistic of anti-tumorigenic effect of TLR3 is well established, wherein TRIF dependent classical pathway induces apoptosis in cancer cells through the endosomal compartment.

However, there are several contradictory reports on the working mechanism and failure in clinical trials. It has also been reported to promote cellular proliferation in head and neck and multiple myeloma cell lines via c-Myc- and NF-κB, respectively following TLR3 ligand poly(I:C) stimulation (Braunstein et al., 2018). In squamous cell carcinomas of the head and neck (HNSCC), triggering the TLR3 signaling pathway along with cisplatin that induces production of the pro-inflammatory cytokine IFN-β, IL-6 and CCL5 to promote cellular survival (Chuang et al., 2018). In a study on metastatic intestinal epithelial cells (IECs), full-length and cleaved form of surface TLR3 has been reported but activation of endosomal TLR3 by poly(I:C) neither induced IFN-β production, nor it induced cell-apoptosis that implies towards the cell surface signaling of TLR3 (Bugge et al., 2017). We had previously reported the surface localization of TLR3 and its proliferative effect on breast cancer cells (Bondhopadhyay et al., 2015). In the present study, we have shown the proliferation of two different types of breast cancer cell lines viz a triple-negative breast cancer cell MDA-MB-231 and an estrogen receptor-positive cells - T47D cells by induction of TLR3 ligand which was downregulated by the addition of the MyD88 inhibitor.

Taken together, the present work generates valuable evidence on the TLR3 mediated alternative signaling of the TLR3-MyD88-IRAK1-TRAF6-TAK1-TAB-NF-κB axis leading to upregulation of IL-6 and cyclinD1 and culminating in proliferation of breast cancer cells; a response that is regulated via MyD88 gateway. Accordingly, a mechanism scheme of alternative TLR3 signal transduction responsible for the proliferation of the cancer cells have been presented (Fig. 7). The outcome of the present study will help in better understanding of the differential response observed in therapeutic use of TLR3. Based on the findings, it is recommended that a targeted delivery of TLR3 ligand to the endosomal compartment, bypassing the MyD88 signaling and subsequently causing activation of TRIF signaling can trigger the apoptotic cascade in cancer cells.

**Fig 7.**
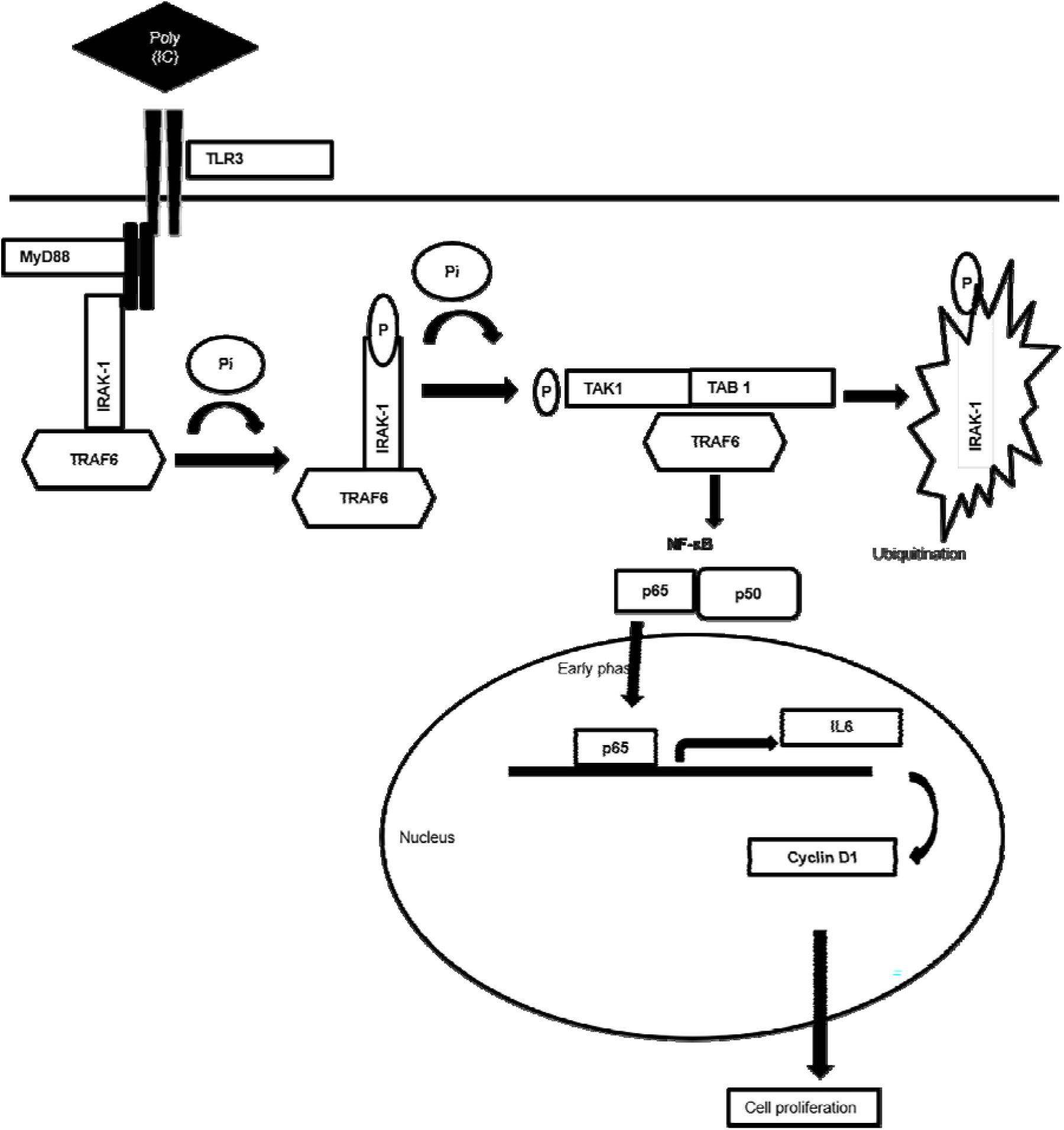
Schematic diagram showing mechanistic of MyD88 adopter mediated surface TLR3 signalling. The diagram illustrating the how TLR3 ligand poly(I:C)-induce the recruitment of MyD88 complex and activation of downstream signaling cascade. Downstream activation of IRAK-1, TAK1, TRAF6 and TAB1 enables translocation of NF-κB, p65 to nucleus to induce the secretion of proinflammatory cytokine IL-6 that induces cell proliferation via cyclin D1.

## The abbreviations used are

ANOVA: analysis of variance;
dsRNA: double-stranded RNA;
ER: endoplasmic reticulum;
HMW: high-molecular-weight;
MyD88: Myeloid differentiation primary response 88;
poly(I:C): Polyinosinic: polycytidylic acid;
TLR 3: Toll-like receptor 3;
TRIF: TIR domain–containing adaptor-inducing interferon.

## Data Availability Statement

The data used to support the findings of this study are available from the corresponding author upon request.

## Ethical approval

Not applicable

## Authors contribution

AS performed the experiments. RSD helped in some experiments. AB designed and Supervise the entire study.

## Acknowledgements

The authors acknowledge DST – SERB for providing funds for the project. The Authors also like to express they’re thanks to CRNN, the University of Calcutta and Prof. Suman Dhar, Special Centre for Molecular Medicine, JNU for providing their Flowcytometry facility for this work. The authors also thank NCCS, Pune for providing the breast cancer cell line from the National facility of cell repository. Authors also acknowledge central imaging facility, Department of Biological Sciences, ISSER Kolkata and CIF, IIT Gandhinagar for providing confocal microscope facility.

## Declaration

[The part of the current work has been presented in ESMO Breast Cancer conference, Berlin 2019 held on May 2-4, 2019 and appeared in Abstract book of ESMO Breast cancer 2019, at Annals of Oncology, Volume 30. Issue supplement _3].

---

**Supplementary Fig 1.**
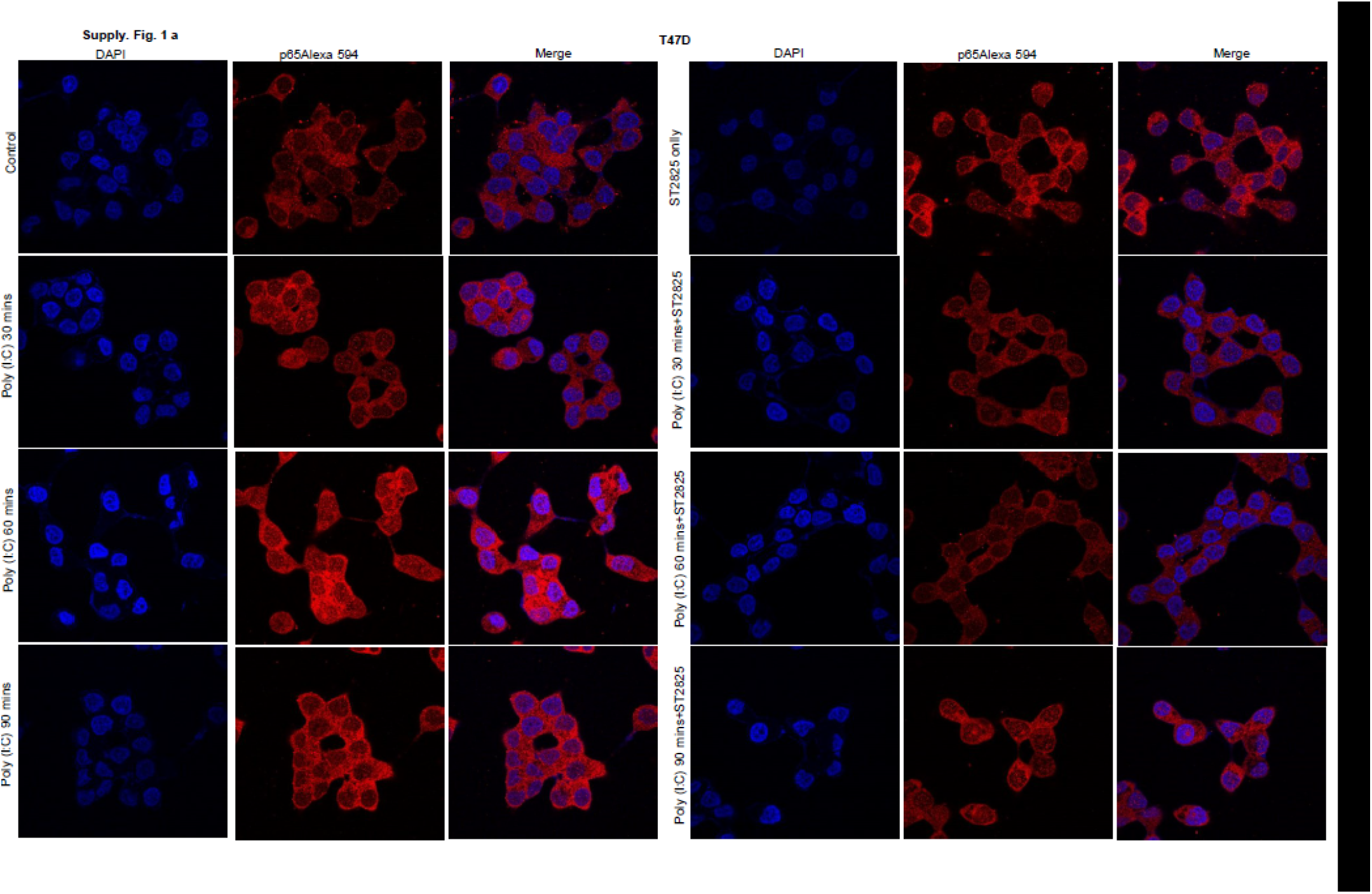
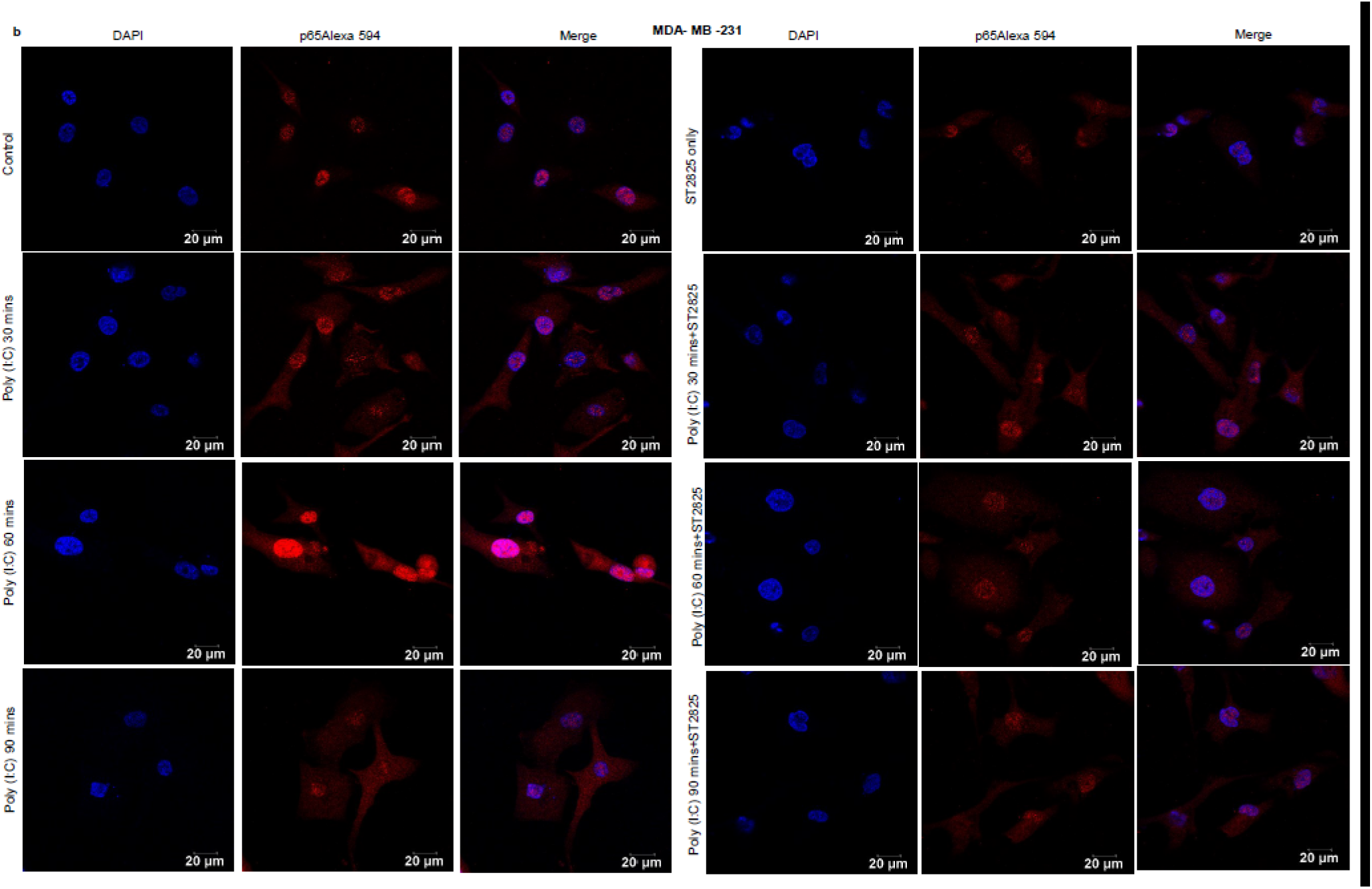
ST2825 attenuates poly(I:C) induced MyD88 dependent early phase activation of NF-κB activation for nuclear translocation of p65 in time dependent manner. **(A)** T47D cells and **(B)** MDA-MB-231 cells were pre-treated with ST2825 for 4 hours prior to addition of poly(10μg/mL) for 30 minutes, 60 minutes and 90 minutes. Cells were stained with antibody against p65 subunit of NF-κB and Alexa 594 tagged secondary antibody and counterstained with DAPI and analyzed acquired through confocal microscope.

## References

1. Alt, J. R., Cleveland, J. L., Hannink, M., & Diehl, J. A. (2000). Phosphorylation-dependent regulation of cyclin D1 nuclear export and cyclin D1–dependent cellular transformation. Genes & Dev. 14, 3102–3114. http://doi.org/10.1101/gad.854900.

2. Aranda, F., Vacchelli, E., Obrist, F., Eggermont, A., Galon, J., Sautès-Fridman, C., et al. (2014). Trial Watch: Toll-like receptor agonists in oncological indications. Oncoimmunology 3, e29179. https://doi.org/10.4161/onci.29179.

3. Bondhopadhyay, B., Moirangthem, A., & Basu, A. (2015). Innate adjuvant receptor Toll-like receptor 3 can promote breast cancer through cell surface. Tumor. Biol. 36, 1261–1271. https://doi.org/10.1007/s13277-014-2737-8.

4. Brasier, A. R. (2010). The nuclear factor-κB—interleukin-6 signalling pathway mediating vascular inflammation. Cardiovasc. Res. 86, 211–218. https://doi.org/10.1093/cvr/cvq076.

5. Braunstein, M. J., Kucharczyk, J., & Adams, S. (2018). Targeting toll-like receptors for cancer therapy. Target. Oncol. 13,583–598. https://doi.org/10.1007/s11523-018-0589-7.

6. Brown, J., Wang, H., Hajishengallis, G. N., & Martin, M. (2011). TLR-signaling networks: an integration of adaptor molecules, kinases, and cross-talk. J. Dent. Res. 90, 417–427. https://doi.org/10.1177/0022034510381264.

7. Bugge, M., Bergstrom, B., Eide, O. K., Solli, H., Kjønstad, I. F., Stenvik, J., et al. (2017). Surface Toll-like receptor 3 expression in metastatic intestinal epithelial cells induces inflammatory cytokine production and promotes invasiveness. J. Biol. Chem. 292, 15408–15425. https://doi.org/10.1074/jbc.M117.784090

8. Chen, J., He, J., Yang, Y., & Jiang, J. (2018). An analysis of the expression and function of myeloid differentiation factor 88 in human osteosarcoma. Oncol. Lett. 16, 4929–4936. https://doi.org/10.3892/ol.2018.9297.

9. Choe, J., Kelker, M. S., & Wilson, I. A. (2005). Crystal structure of human toll-like receptor 3 (TLR3) ectodomain. Science 309, 581–585. https://doi.org/10.1126/science.1115253

10. Chuang, H. C., Chou, M. H., Chien, C. Y., Chuang, J. H., & Liu, Y. L. (2018). Triggering TLR3 pathway promotes tumor growth and cisplatin resistance in head and neck cancer cells. Oral. Oncol. 86, 141–149. https://doi.org/10.1016/j.oraloncology.2018.09.015.

11. Conforti, R., Ma, Y., Morel, Y., Paturel, C., Terme, M., Viaud, S., et al. (2010). Opposing effects of toll-like receptor (TLR3) signaling in tumors can be therapeutically uncoupled to optimize the anticancer efficacy of TLR3 ligands. Cancer. Res. 70, 490–500. https://doi.org/10.1158/0008-5472.CAN-09-1890.

12. Cui, W., Xiao, N., Xiao, H., Zhou, H., Yu, M., Gu, J., et al. (2012). β-TrCP-mediated IRAK1 degradation releases TAK1-TRAF6 from the membrane to the cytosol for TAK1-dependent NF-κB activation. Mol. Cell. Biol. 32,3990–4000. https://doi.org/10.1128/MCB.00722-12.

13. Deng, Y., Sun, J., & Zhang, L. D. (2016). Effect of ST2825 on the proliferation and apoptosis of human hepatocellular carcinoma cells. Genet. Mol. Res. 15,15016826. https://doi.org/10.4238/gmr.15016826.

14. Dong, W., Liu, Y., Peng, J., Chen, L., Zou, T., Xiao, H., et al. (2006). The IRAK-1-BCL10-MALT1-TRAF6-TAK1 cascade mediates signaling to NF-κB from Toll-like receptor 4. J. Biol. Chem. 281,26029–26040. https://doi.org/10.1074/jbc.M513057200.

15. Gambara, G., Desideri, M., Stoppacciaro, A., Padula, F., De Cesaris, P., Starace, D., et al. (2015). TLR 3 engagement induces IRF□3□dependent apoptosis in androgen□sensitive prostate cancer cells and inhibits tumour growth in vivo. J. Cell. Mol. Med. 19,327–339. https://doi.org/10.1111/jcmm.12379.

16. González-Reyes, S., Marín, L., González, L., González, L. O., del Casar, J. M., Lamelas, M. L., et al. (2010). Study of TLR3, TLR4 and TLR9 in breast carcinomas and their association with metastasis. BMC cancer 60, 217–226. https://doi.org/10.1186/1471-2407-10-665.

17. Han, S. J., Ko, H. M., Choi, J. H., Seo, K. H., Lee, H. S., Choi, E. K., et al. (2002). Molecular mechanisms for lipopolysaccharide-induced biphasic activation of nuclear factor-κB (NF-κB). J. Biol. Chem. 277, 44715–44721. https://doi.org/10.1074/jbc.M202524200.

18. Helminen, O., Huhta, H., Lehenkari, P. P., Saarnio, J., Karttunen, T. J., & Kauppila, J. H. (2016). Nucleic acid-sensing toll-like receptors 3, 7 and 8 in esophageal epithelium, barrett’s esophagus, dysplasia and adenocarcinoma. Oncoimmunology 5, e1127495. https://doi.org/10.1080/2162402X.2015.1127495.

19. Ho, V., Lim, T. S., Lee, J., Steinberg, J., Szmyd, R., Tham, M., et al. (2015). TLR3 agonist and Sorafenib combinatorial therapy promotes immune activation and controls hepatocellular carcinoma progression. Oncotarget 6, 27252. https://doi.org/10.18632/oncotarget.4583.

20. Hsiao, H. M., Thatcher, T. H., Levy, E. P., Fulton, R. A., Owens, K. M., Phipps, R. P., et al. (2014). Resolvin D1 Attenuates Polyinosinic-Polycytidylic Acid–Induced Inflammatory Signaling in Human Airway Epithelial Cells via TAK1. J. Immunol. 193,4980–4987. https://doi.org/10.4049/jimmunol.1400313.

21. Jensen, L. E., & Whitehead, A. S. (2003). Pellino3, a novel member of the Pellino protein family, promotes activation of c-Jun and Elk-1 and may act as a scaffolding protein. J. Immunol. 171, 1500–1506. https://doi.org/10.4049/jimmunol.171.3.1500.

22. Jia, D., & Wang, L. (2015). The other face of TLR3: A driving force of breast cancer stem cells. Mol. Cell. Oncol. 2, e981443. https://doi.org/10.4161/23723556.2014.981443.

23. Jia, D., Yang, W., Li, L., Liu, H., Tan, Y., Ooi, S., et al. (2015). β-Catenin and NF-κ B coactivation triggered by TLR3 stimulation facilitates stem cell-like phenotypes in breast cancer. Cell. Death. Differ. 22, 298–310. https://doi.org/10.1038/cdd.2014.145.

24. Johnsen, I. B., Nguyen, T. T., Ringdal, M., Tryggestad, A. M., Bakke, O., Lien, E., et al. (2006). Toll□like receptor 3 associates with c□Src tyrosine kinase on endosomes to initiate antiviral signaling. EMBO J. 25, 3335–3346. https://doi.org/10.1038/sj.emboj.7601222.

25. Kawasaki, T., & Kawai, T. (2014). Toll-like receptor signaling pathways. Front. Immunol. 5, 461. https://doi.org/10.3389/fimmu.2014.00461.

26. Klein, E. A., & Assoian, R. K. (2008). Transcriptional regulation of the cyclin D1 gene at a glance. J. Cell. Sci. 121,3853–3857. https://doi.org/10.1242/jcs.039131.

27. Kong, F., Liu, Z., Jain, V. G., Shima, K., Suzuki, T., Muglia, L. J., et al. (2017). Inhibition of IRAK1 ubiquitination determines glucocorticoid sensitivity for TLR9-induced inflammation in macrophages. J. Immunol. 199, 3654–3667. https://doi.org/10.4049/jimmunol.1700443.

28. Loiarro, M., Capolunghi, F., Fanto, N., Gallo, G., Campo, S., Arseni, B., et al. (2007). Pivotal Advance: Inhibition of MyD88 dimerization and recruitment of IRAK1 and IRAK4 by a novel peptidomimetic compound. J. Leukocyte. Biol. 82, 801–810, https://doi.org/10.1189/jlb.1206746.

29. Loiarro, M., Ruggiero, V., & Sette, C. (2010). Targeting TLR/IL-1R signalling in human diseases. Mediat. Inflamm. 2010, 1–12. https://doi.org/10.1155/2010/674363.

30. Ma, L., Feng, L., Ding, X., & Li, Y. (2018). Effect of TLR4 on the growth of SiHa human cervical cancer cells via the MyD88-TRAF6-TAK1 and NF-κB-cyclin D1-STAT3 signaling pathways. Oncol. Lett. 15, 3965–3970. https://doi.org/10.3892/ol.2018.7801.

31. Matijevic Glavan, T., & Pavelic, J. (2014). The exploitation of Toll-like receptor 3 signaling in cancer therapy. Curr. Pharm. Des. 20, 6555–6564. https://doi.org/10.2174/1381612820666140826153347.

32. McFarland, B. C., Hong, S. W., Rajbhandari, R., Twitty Jr, G. B., Gray, G. K., Yu, H., et al. (2013). NF-κB-induced IL-6 ensures STAT3 activation and tumor aggressiveness in glioblastoma. PLoS ONE 8, e78728. https://doi.org/10.1371/journal.pone.0078728.

33. Noursadeghi, M., Tsang, J., Haustein, T., Miller, R. F., Chain, B. M., & Katz, D. R. (2008). Quantitative imaging assay for NF-κB nuclear translocation in primary human macrophages. J. Immunol. Methods. 329,194–200. https://doi.org/10.1016/j.jim.2007.10.015.

34. O’Neill, L. A., & Bowie, A. G. (2007). The family of five: TIR-domain-containing adaptors in Toll-like receptor signalling. Nat. Rev. Immunol. 7, 353–364. https://doi.org/10.1038/nri2079.

35. Oshiumi, H., Matsumoto, M., Funami, K., Akazawa, T., & Seya, T. (2003). TICAM-1, an adaptor molecule that participates in Toll-like receptor 3–mediated interferon-β induction. Nat. Immunol. 4, 161–167. https://doi.org/10.1038/ni886.

36. Rhyasen, G. W., & Starczynowski, D. T. (2015). IRAK signalling in cancer. Br. J. Cancer. 112, 232–237. https://doi.org/10.1038/bjc.2014.513.

37. Salaun, B., Coste, I., Rissoan, M. C., Lebecque, S. J., & Renno, T. (2006). TLR3 can directly trigger apoptosis in human cancer cells. J. Immunol. 176, 4894–4901. https://doi.org/10.4049/jimmunol.176.8.4894.

38. Salaun, B., Zitvogel, L., Asselin-Paturel, C., Morel, Y., Chemin, K., Dubois, C., et al. (2011). TLR3 as a biomarker for the therapeutic efficacy of double-stranded RNA in breast cancer. Cancer Res. 71, 1607–1614. https://doi.org/10.1158/0008-5472.CAN-10-3490.

39. Schau, I., Michen, S., Hagstotz, A., Janke, A., Schackert, G., Appelhans, D., et al. (2019). Targeted delivery of TLR3 agonist to tumor cells with single chain antibody fragment-conjugated nanoparticles induces type I-interferon response and apoptosis. Sci. Rep. 9, 3299. https://doi.org/10.1038/s41598-019-40032-8.

40. Shiratori, E., Itoh, M., & Tohda, S. (2017). MYD88 inhibitor ST2825 suppresses the growth of lymphoma and leukaemia cells. Anticancer Res. 37, 6203–6209. https://doi.org/10.21873/anticanres.12070.

41. Smith, M., García-Martínez, E., Pitter, M. R., Fucikova, J., Spisek, R., Zitvogel, L., et al. (2018). Trial Watch: Toll-like receptor agonists in cancer immunotherapy. Oncoimmunology 7, e1526250. https://doi.org/10.1080/2162402X.2018.1526250.

42. Takeda, K., & Akira, S. (2004). TLR signaling pathways. Semin. Immunol. 16, 3–9. https://doi.org/10.1016/j.smim.2003.10.003.

43. Torres-Luquis, O. J., & Mohammed, S. (2019). Abstract 2019A: TLR3 facilitate breast cancer metastasis to lymph node. Atlanta, GA. Philadelphia (PA): AACR; Cancer Res 2019; 79(13 Suppl): Abstract nr 2019A. https://doi.org/10.1158/1538-7445.AM2019-2019A.

44. Xiong, Y., Qiu, F., Piao, W., Song, C., Wahl, L. M., & Medvedev, A. E. (2011). Endotoxin tolerance impairs IL-1 receptor-associated kinase (IRAK) 4 and TGF-β-activated kinase 1 activation, K63-linked polyubiquitination and assembly of IRAK1, TNF receptor-associated factor 6, and IκB kinase γ and increases A20 expression. J. Biol. Chem. 286,7905–7916. https://doi.org/10.1074/jbc.M110.182873.

45. Yamamoto, M., Sato, S., Hemmi, H., Hoshino, K., Kaisho, T., Sanjo, H., et al. (2003). Role of adaptor TRIF in the MyD88-independent toll-like receptor signaling pathway. Science 301, 640–643. https://doi.org/10.1126/science.1087262.

46. Yang, H., Qi, H., Ren, J., Cui, J., Li, Z., Waldum, H. L., & Cui, G. (2014). Involvement of NF-κB/IL-6 pathway in the processing of colorectal carcinogenesis in colitis mice. Int. J. Inflam. 2014,1–7. https://doi.org/10.1155/2014/130981.

